# Identifying Novel Therapeutic Targets for Overcoming TNBC Chemo Resistance Through Comprehensive CRISPR-Cas9 Genome Screening

**DOI:** 10.1101/2024.05.14.594192

**Authors:** Shuai Shao, Shangjia Li, Shan Tang, Kunjie Fan, Lang Li

## Abstract

Triple-negative breast cancer (TNBC) represents 15-20% of cases but disproportionately contributes to 35% of breast cancer deaths. Chemotherapy resistance remains a significant challenge in TNBC treatment. In this study, we identified the MDA-MB-231 cell line as the most representative model for TNBC chemotherapy-poor responders by comparing genomic profiles from TNBC cell lines and patient samples. We performed a genome-wide CRISPR-Cas9 screen and RNAseq analysis in MDA-MB-231 cells to uncover potential synthetic lethal targets for cisplatin/doxorubicin treatment.

Our analysis confirmed the involvement of known essential genes in DNA damage repair and regulation of DNA replication pathways, such as BCL2L1, ATM, CDC25B, and NBN, in sensitizing cells to cisplatin/doxorubicin. Additionally, We identified hundreds of previously unrecognized genes and pathways related to DNA repair, G2/M DNA damage checkpoint, AMPK signaling, and mTOR signaling. The observed differences between transcriptomic responses and essential pathways from the CRISPR screen suggest a complex regulatory system in cellular response to DNA damage drugs. By combining various data analysis methods and biological experimental approaches, we have pinpointed several promising genes, such as MCM9 and NEPPS, which could serve as potential drug targets to overcome chemoresistance.

Overall, our approach efficiently identified essential genes with potential synthetic lethal interactions with cisplatin/doxorubicin, offering new possibilities for combination therapies in chemo resistant TNBC patients.

## INTRODUCTION

Triple-negative breast cancer (TNBC) represents a breast cancer subtype distinguished by the absence of three pivotal receptors: estrogen receptor (ER), progesterone receptor (PR), and human epidermal growth factor receptor 2 (HER2) ^1^. These receptors substantially influence the proliferation and dissemination of breast cancer cells. In most breast cancer cases, therapeutic interventions target one or more of these receptors to impede or halt cancer progression^2^. Nevertheless, the lack of these receptors in TNBC renders it a particularly formidable challenge for treatment^3^.

A combination of surgical intervention, radiotherapy, and chemotherapy constitutes the conventional treatment regimen for TNBC ^4^. Cisplatin and doxorubicin, two frontline chemotherapeutic agents, are employed in managing triple-negative breast cancer. Cisplatin, a platinum-based chemotherapy compound, functions by forming covalent bonds with DNA, potentially resulting in DNA damage and subsequent cellular demise ^5,6^. Doxorubicin, another DNA-damaging drug, is an anthracycline chemotherapeutic agent that primarily operates by inflicting damage upon the DNA of cancer cells. Its mechanism encompasses DNA double helix intercalation, topoisomerase II inhibition, and free radical generation, culminating in DNA damage and eventual cell death ^7^. While these chemotherapeutic drugs effectively eradicate tumor cells, resistance has been observed in TNBC patients ^8,9^. In large clinical trials, it has been observed that approximately half of the patients with triple-negative breast cancer (TNBC) have residual cancer after undergoing neoadjuvant chemotherapy (NACT) ^10,11^. Furthermore, around 40% of residual disease patients will eventually develop distant metastasis ^12^. These findings underscore the importance of identifying more effective treatment strategies for TNBC poor chemo drug responders.

In recent years, advances in understanding TNBC’s molecular characteristics have led to new therapeutic options. Immunotherapy, such as atezolizumab^13^, has shown promise when combined with nab-paclitaxel for advanced TNBC. PARP inhibitors (e.g., olaparib, niraparib, talazoparib) ^14^ ^15^ have emerged as valuable, particularly when combined with chemotherapy agents for metastatic TNBC patients. Antibody-drug conjugates, like sacituzumab govitecan^16^, selectively deliver cytotoxic agents to cancer cells, minimizing damage to healthy cells. Moreover, targeted therapies, including PI3K and mTOR inhibitors, offer new TNBC treatment avenues^17^. These advances underscore the potential for novel, personalized therapies based on TNBC’s molecular landscape.

These emerging treatments capitalize on the concept of synergistic lethality, which occurs when the simultaneous inhibition or disruption of two or more genes or pathways results in cell death ^18^.The combination of PARP inhibitors and HRR deficiency successfully applies this concept in cancer therapy ^19^. PARP inhibitors are promising as combined drug therapy with DNA-damaging drugs for treating BRCA1 or BRCA2 mutated, HER2-negative breast cancers. Olaparib and talazoparib have received FDA approval for this purpose ^20^. Poly (ADP-ribose) polymerase (PARP) is an enzyme related with DNA repair, explicitly repairing single-strand DNA breaks through the base excision repair (BER) pathway ^21^. BRCA1 and BRCA2 genes play an important role in another type of DNA repair called homologous recombination repair (HRR) ^22^. BRCA1 or BRCA2 mutations cells rely on alternative repair pathways like BER to maintain genomic integrity. Inhibiting PARP leads to unrepaired single-strand breaks converting into double-strand during replication. Without functional HRR, cells cannot repair these breaks, resulting in genomic instability and cell death^23^.

The CRISPR/Cas9 system is a versatile and powerful gene editing tool that disrupts or modifies specific genes ^24,25^. Genome-wide CRISPR screening allows researchers to systematically examine the entire genome to identify genes whose knockout or modification can lead to particular phenotypes, such as increased sensitivity to chemotherapy or synthetic lethality combined with another genetic disruption^26,27^ ^28,29^. In our study, we conduct a genome-wide CRISPR screening using the TKOV3 library to identify potential druggable targets that could help overcome resistance to DNA-damaging chemotherapeutic agents, such as cisplatin and doxorubicin, in TNBC patients. Our approach focuses on investigating synthetic lethality and potential drug combinations to enhance treatment efficacy and counteract resistance mechanisms.

Before carrying out our CRISPR screening experiment, it is crucial to select an appropriate and representative cell line model for the study. The similarity between the cell model and patients with drug-resistant tumors directly determines whether the drug targets we identify can genuinely progress from preclinical to clinical stages. In Chapter 2, 21 TNBC cell lines from the CCLE database are evaluated by comparing their genomic profiles with those of TNBC poor responders. Among these cell lines, MDA-MB-231 was selected as one of the most representative models for conducting a genome-wide CRISPR screening.

## RESULT

### Pooled genome-wide CRISPR screening on MDA-MB-231 cell lines

We performed a genome-wide CRISPR/cas9 pooled screen to identify genes related to cisplatin or doxorubicin resistance in the most representative cell line, MDA-MB-231. TKOv3sgRNA library, which contains 7,0948 sgRNA targeting 18053 genes, was used for this screening. After lentiviral pooled sgRNA virus and puromycin selection with lower MOI (0.3), the baseline sample was harvested. Additionally, on day 28, samples after cisplatin or doxorubicin, or DMSO treatment with three triplicates were collected (Figure 1). After deep-depth sequencing, the MAGeCKFlute algorithm was used for sequence data analysis. The number of missed genes for all samples is from 200∼400 sgRNA among a total of 7,0948 sgRNA (Figure 2A). With passage time increase, the Gini index was increased from 0.04 to 0.06, which is reasonable because longer treatment will cause the unevenness in pooled sgRNA. From the 3D PCA plot (Figure 2C) and pairwise sample correlation plot (Figure 2D), samples in the same treatment group show a clear pattern and higher correlation value, which indicates that triplicates samples have a constant result. Overall, the data analysis result indicated that our approach successfully produced a CRISPR-based screening of one MDA-MB-231 cell line.

**Figure 1.**
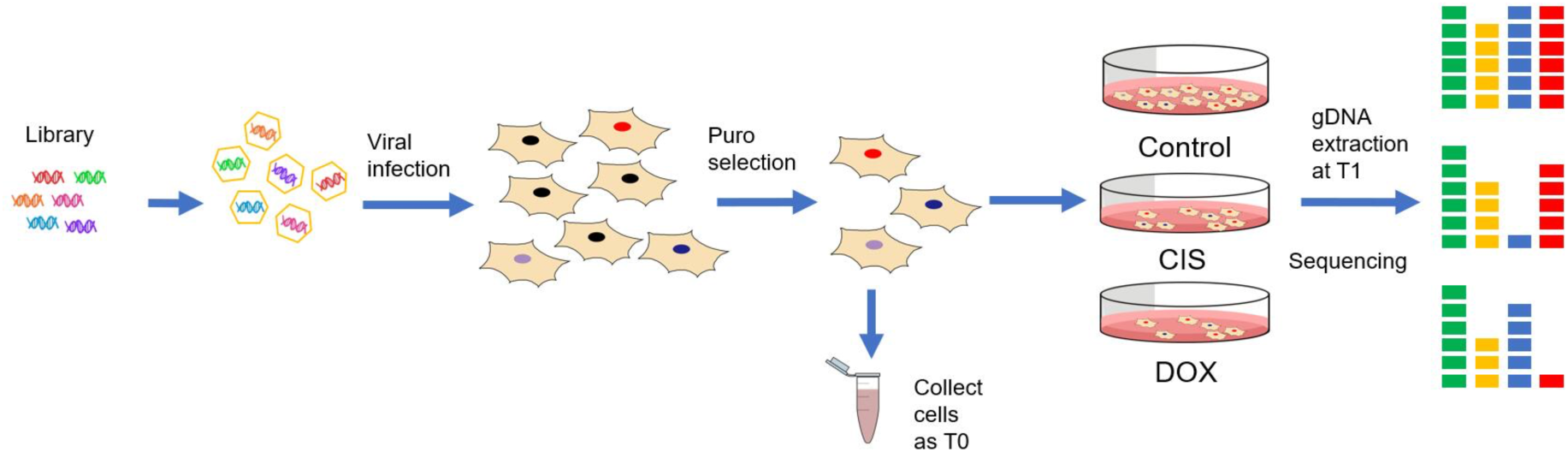
A schematic diagram for genome wide CRISPR by the TKVO3 library.

**Figure 2.**
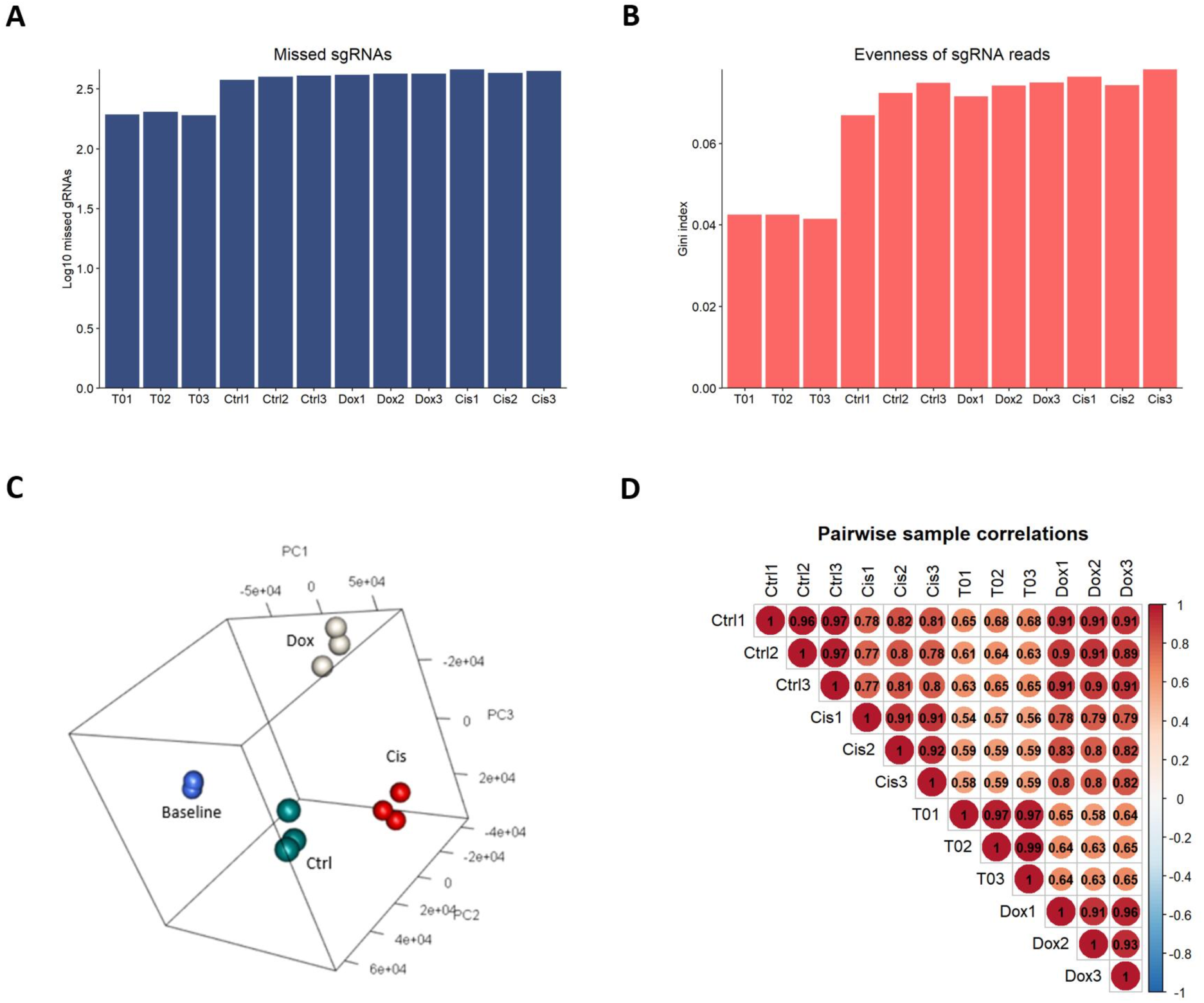
Genome-wide CRISPR-Cas9 negative screens in the MDA-MB-231 cell line. (A) The missed sgRNAs were tested on days 0 (T0), and days 28. (B) The Gini index of sgRNAs on days 0 (T0), and days 28. (C) 3D PCA plot of baseline, control, and two drug treatment groups. PC1 to PC3 can explain more than 80% of total information. (D) Correlation plot between baseline, control, and two drug treatment group, each group containing triplicate samples.

### Essential gene identification and pathway enrichment analysis

Using the MAGeCK MLE algorithm, gene essentiality scores (β scores) were calculated correspondingly in each group. Lower β scores significantly depleted sgRNAs in the final time point. The distribution of normalized sgRNA β scores (Figure 3A for CIS, 6A for DOX) for each group is similar, which indicates the good quality of our negative screening. The square plot for cisplatin treatment (Figure 3C) and doxorubicin treatment (Figure 4C) was generated, the which 96 essential genes (Table 3) and 93 essential genes (Table 4) were identified in cisplatin and doxorubicin correspondingly. KEGG pathways analysis are performed by using essential genes list detected on cisplatin treatment and doxorubicin treatment correspondingly, top 15 highly enriched pathways for each analysis are visualized (Figure 3D and Figure 3D). In both drug treatment studies, the enriched pathways are highly associated with DNA damage/DNA repair signaling pathways.

**Figure 3.**
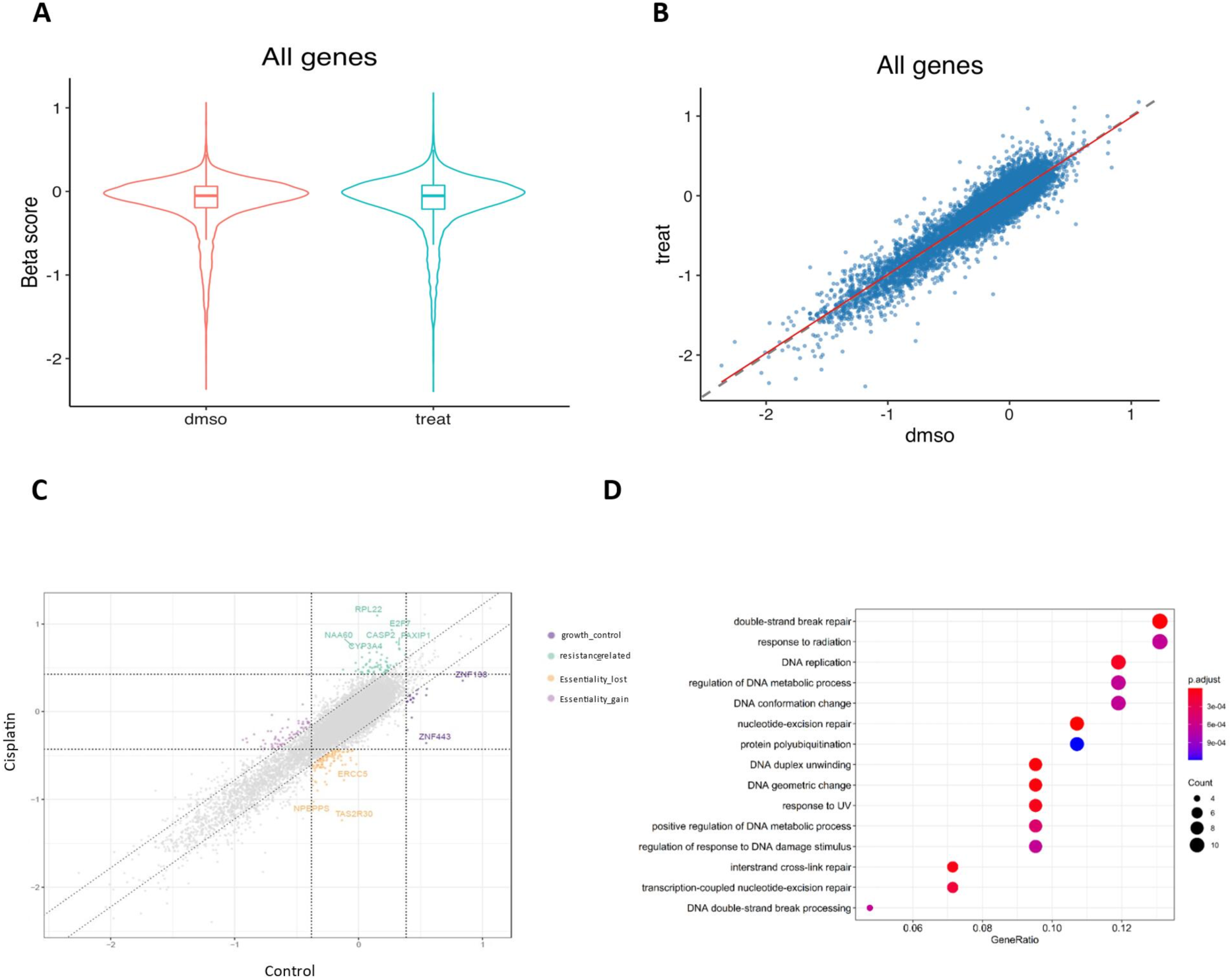
Genome-wide CRISPR knockout screen in TNBC with Cisplatin treatment. (A) Box plot for sgRNA β scores for each group. (B) Normalized distribution of β scores for each group. (C) The gene essentiality scores reported from MAGeCK-MLE in cisplatin treatment. (D) Essential gene enrichment pathways of by KEGG analysis in cisplatin treatment.

**Figure 4.**
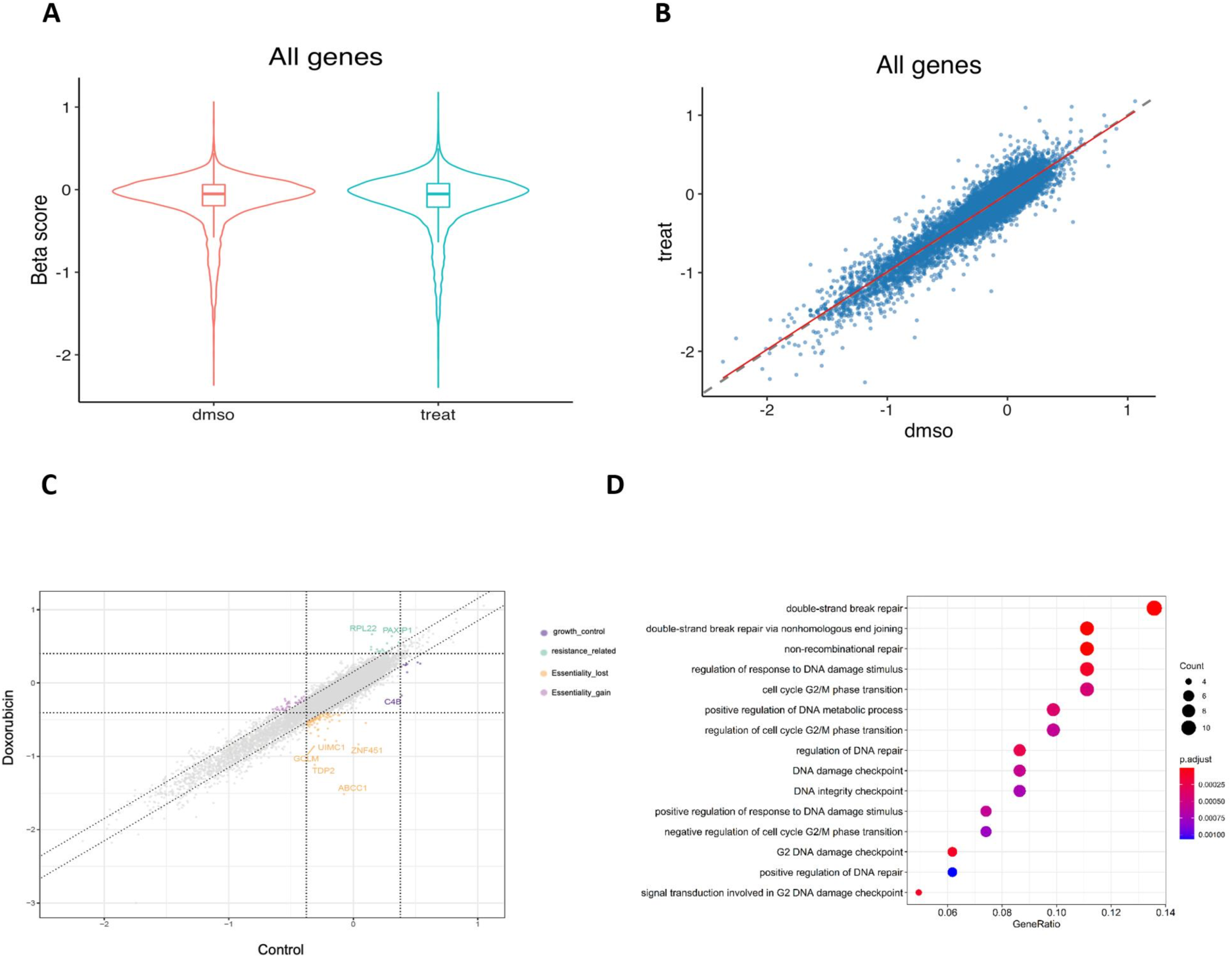
Genome-wide CRISPR knockout screen in TNBC with doxorubicin treatment (A) Box plot for sgRNA β scores for each group. (B) Normalized distribution of β scores for each group. (C) The gene essentiality scores reported from MAGeCK-MLE in cisplatin treatment. (D) Essential gene enrichment pathways of by KEGG analysis in doxorubicin treatment.

**Table 1.**
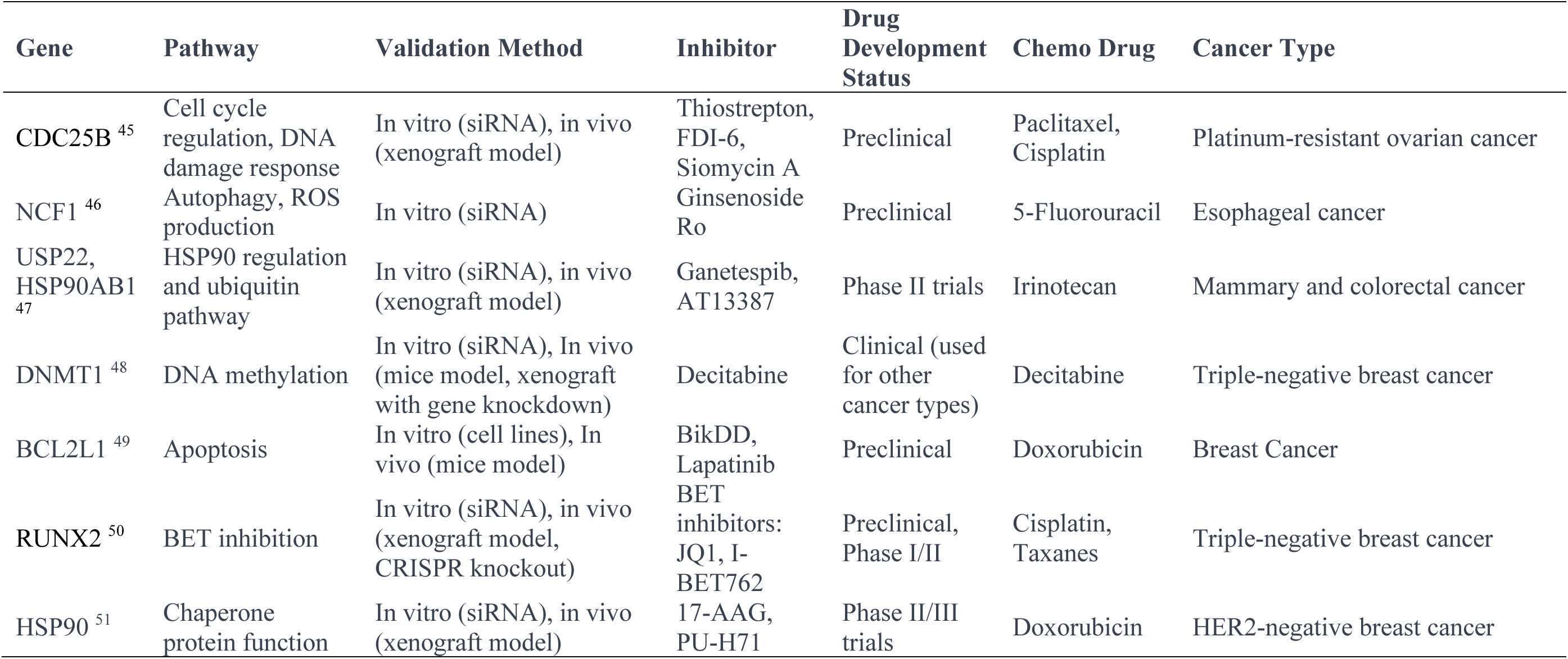
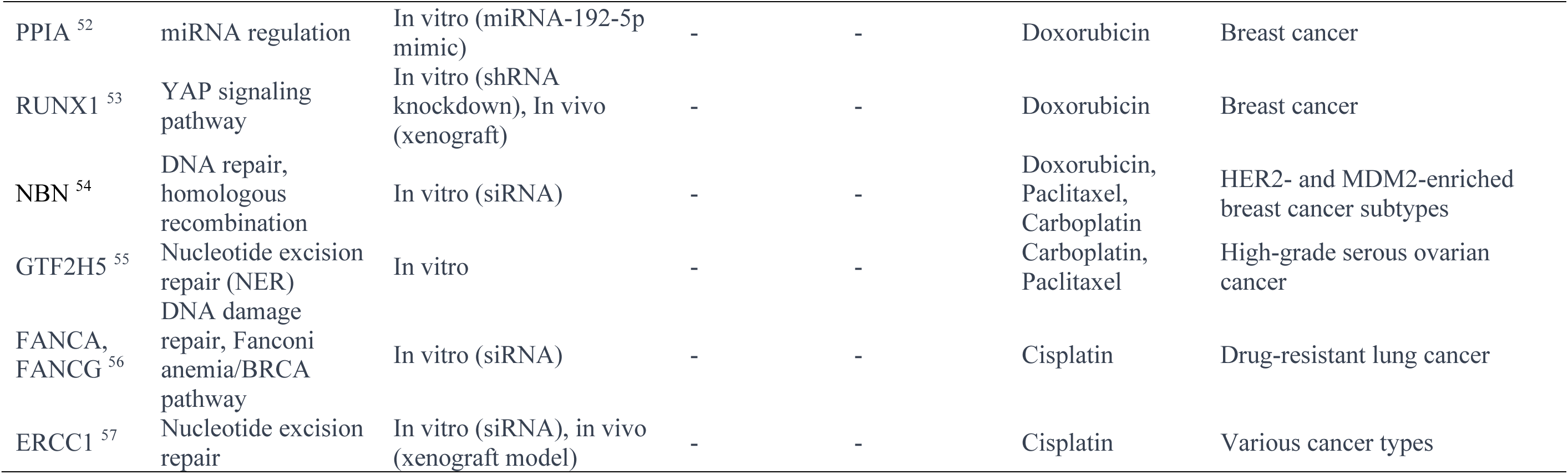
Well studied gene in essential gene list for cisplatin treatment.

**Table 2.**
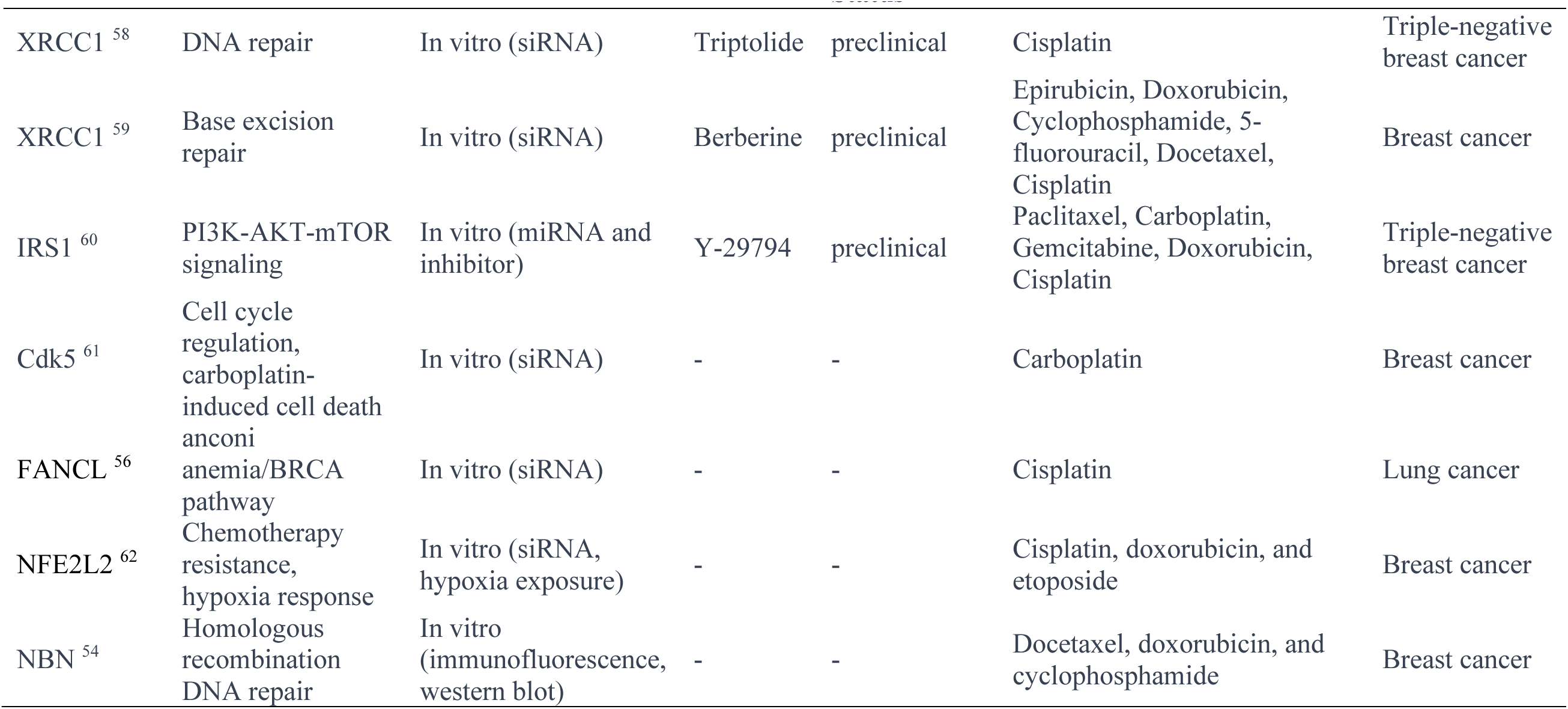
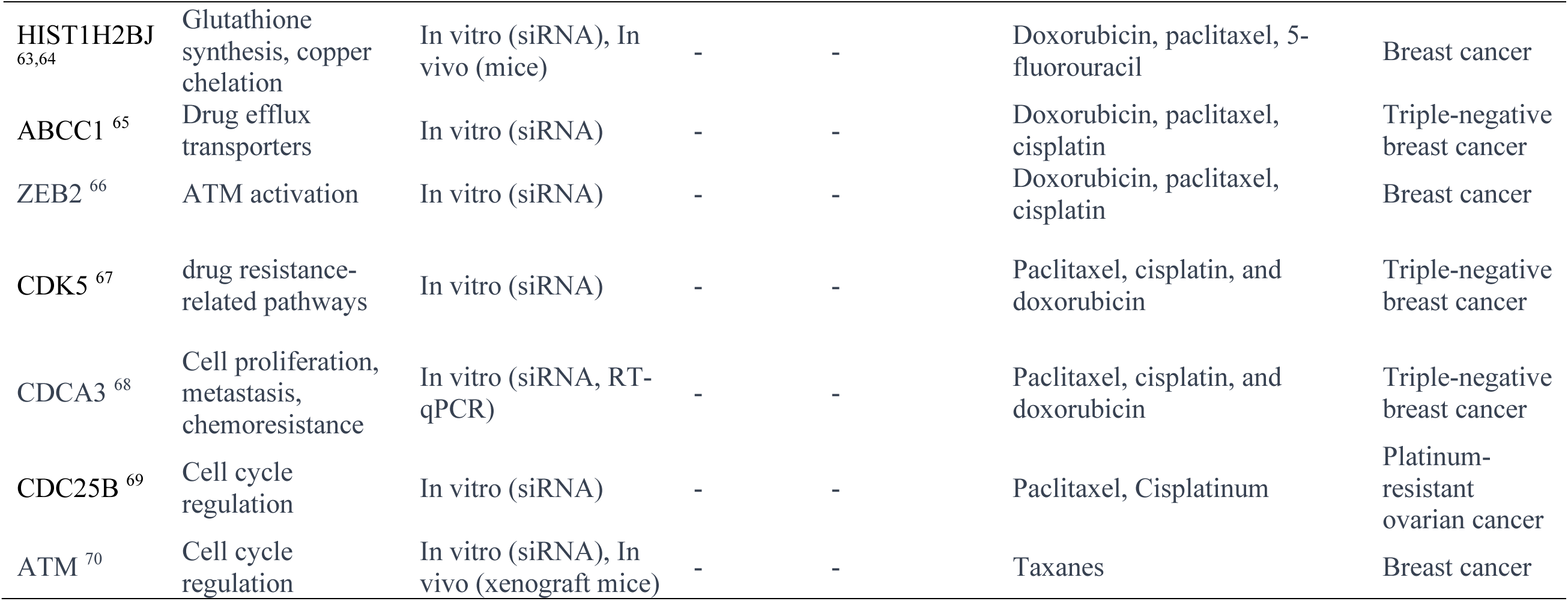
Well studied gene in essential gene list for doxorubicin treatment.

**Table 3.**
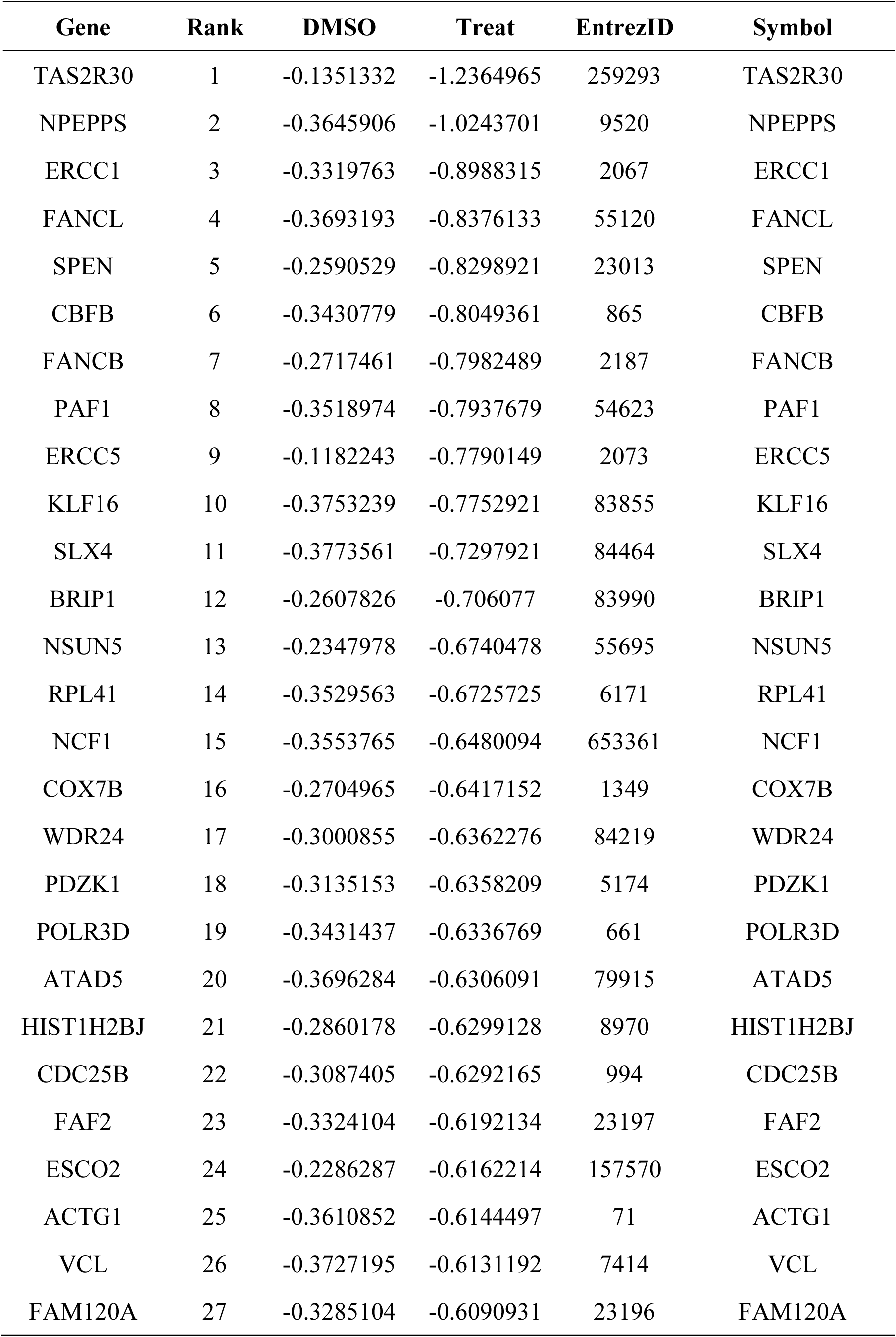
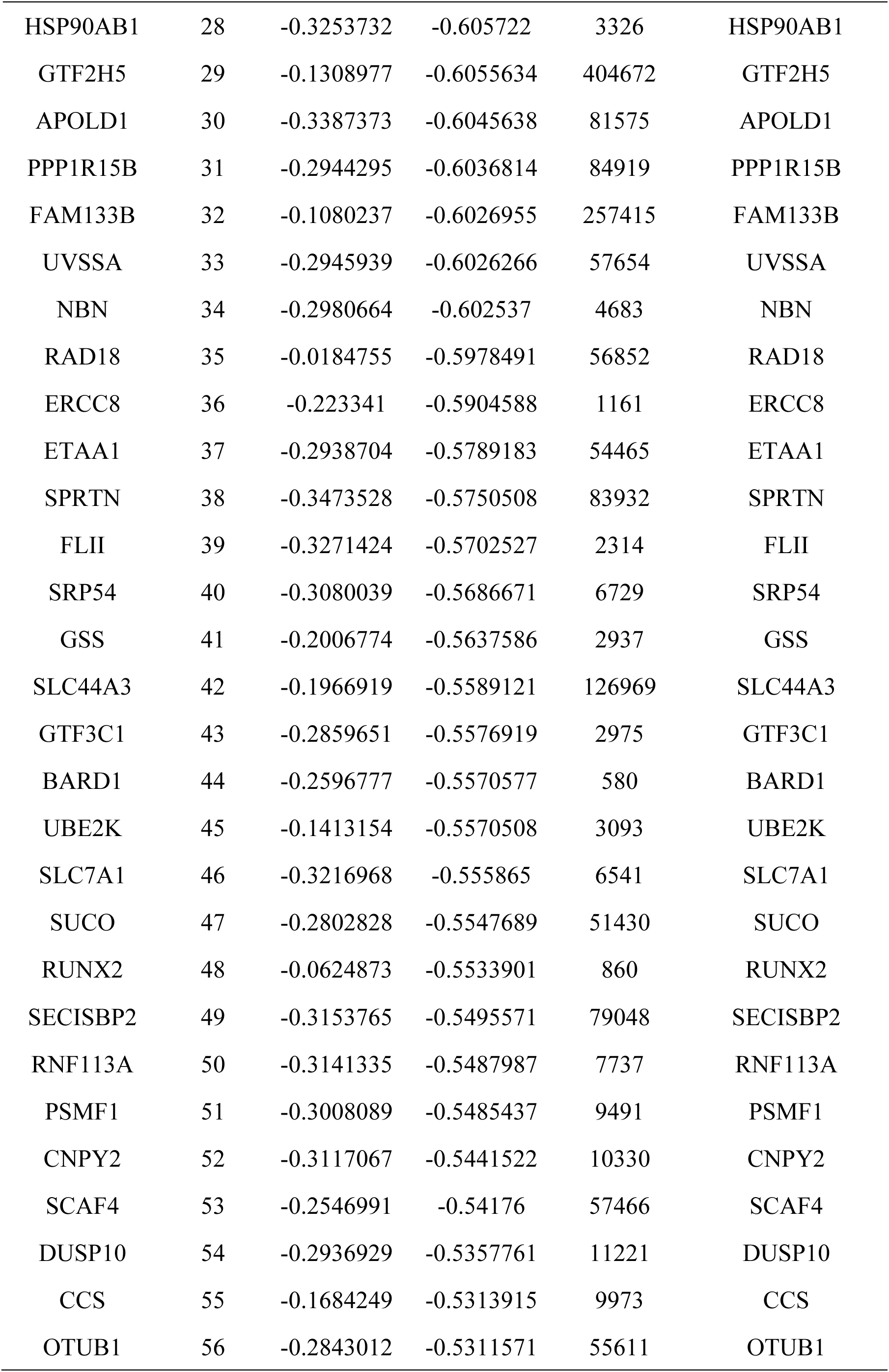
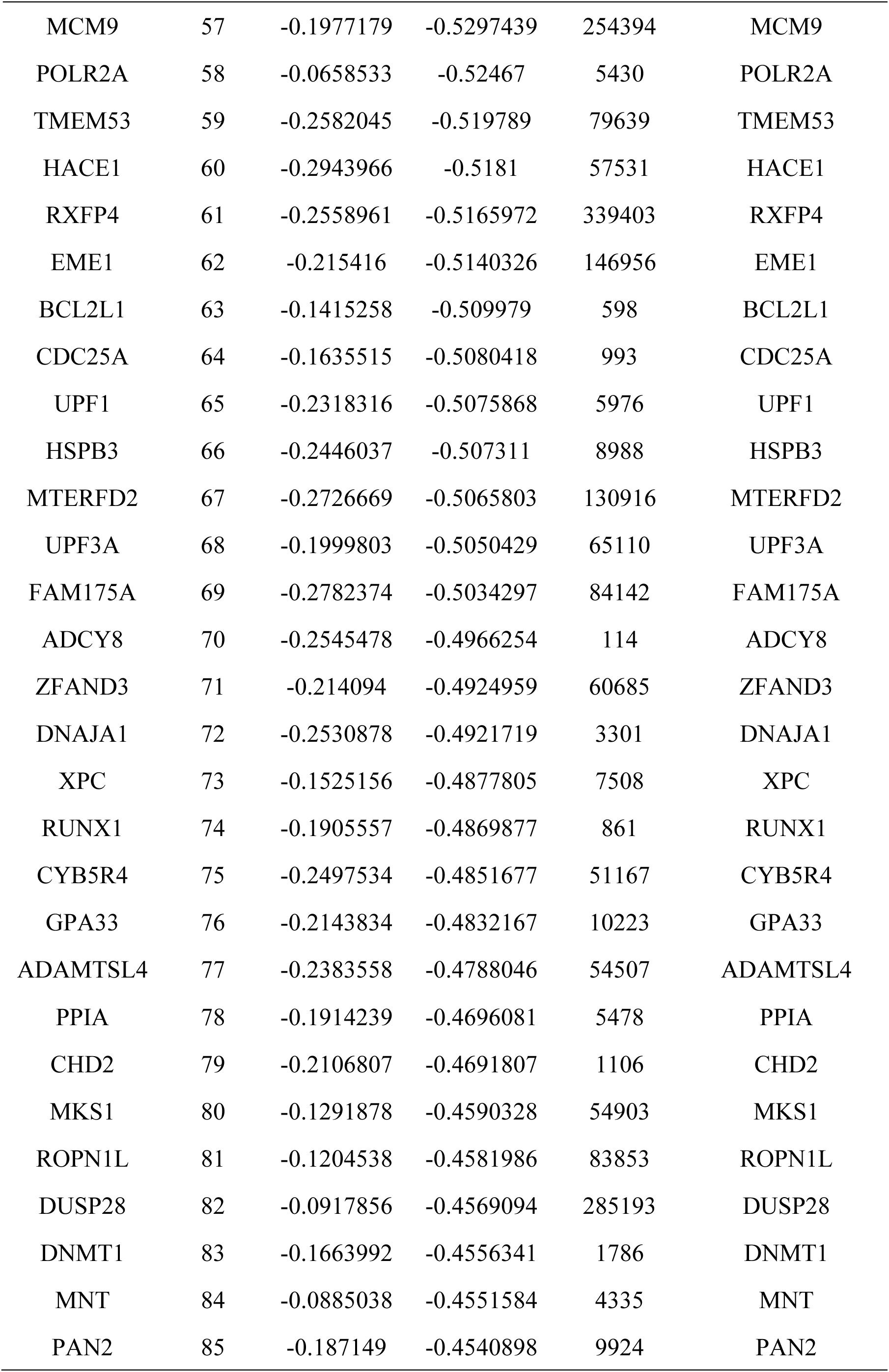
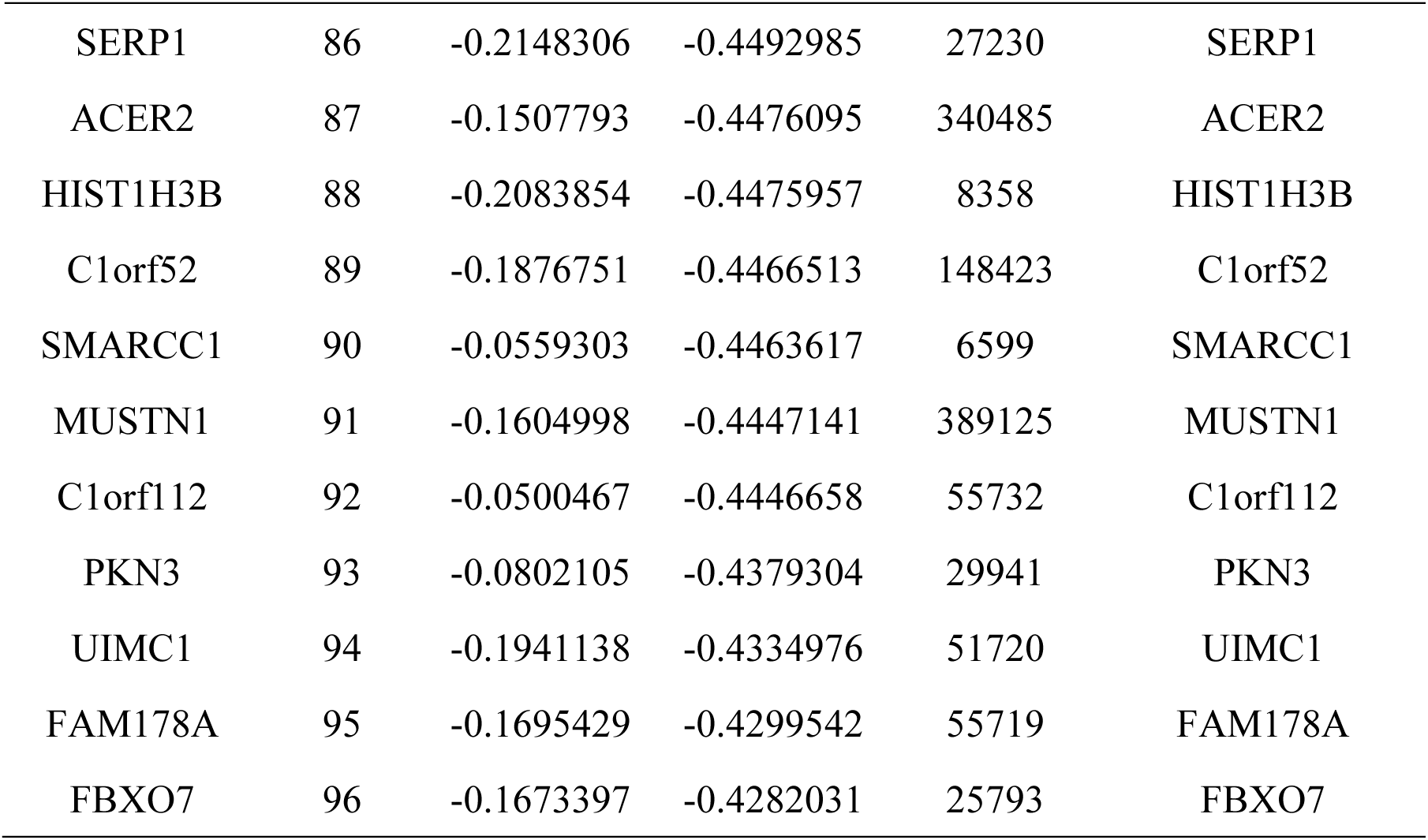
CRISPR essential gene list for Cisplatin treatment.

| **Gene** | **Rank** | **DMSO** | **Treat** | **EntrezID** | **Symbol** |
| --- | --- | --- | --- | --- | --- |
| TAS2R30 | 1 | -0.1351332 | -1.2364965 | 259293 | TAS2R30 |
| NPEPPS | 2 | -0.3645906 | -1.0243701 | 9520 | NPEPPS |
| ERCC1 | 3 | -0.3319763 | -0.8988315 | 2067 | ERCC1 |
| FANCL | 4 | -0.3693193 | -0.8376133 | 55120 | FANCL |
| SPEN | 5 | -0.2590529 | -0.8298921 | 23013 | SPEN |
| CBFB | 6 | -0.3430779 | -0.8049361 | 865 | CBFB |
| FANCB | 7 | -0.2717461 | -0.7982489 | 2187 | FANCB |
| PAF1 | 8 | -0.3518974 | -0.7937679 | 54623 | PAF1 |
| ERCC5 | 9 | -0.1182243 | -0.7790149 | 2073 | ERCC5 |
| KLF16 | 10 | -0.3753239 | -0.7752921 | 83855 | KLF16 |
| SLX4 | 11 | -0.3773561 | -0.7297921 | 84464 | SLX4 |
| BRIP1 | 12 | -0.2607826 | -0.706077 | 83990 | BRIP1 |
| NSUN5 | 13 | -0.2347978 | -0.6740478 | 55695 | NSUN5 |
| RPL41 | 14 | -0.3529563 | -0.6725725 | 6171 | RPL41 |
| NCF1 | 15 | -0.3553765 | -0.6480094 | 653361 | NCF1 |
| COX7B | 16 | -0.2704965 | -0.6417152 | 1349 | COX7B |
| WDR24 | 17 | -0.3000855 | -0.6362276 | 84219 | WDR24 |
| PDZK1 | 18 | -0.3135153 | -0.6358209 | 5174 | PDZK1 |
| POLR3D | 19 | -0.3431437 | -0.6336769 | 661 | POLR3D |
| ATAD5 | 20 | -0.3696284 | -0.6306091 | 79915 | ATAD5 |
| HIST1H2BJ | 21 | -0.2860178 | -0.6299128 | 8970 | HIST1H2BJ |
| CDC25B | 22 | -0.3087405 | -0.6292165 | 994 | CDC25B |
| FAF2 | 23 | -0.3324104 | -0.6192134 | 23197 | FAF2 |
| ESCO2 | 24 | -0.2286287 | -0.6162214 | 157570 | ESCO2 |
| ACTG1 | 25 | -0.3610852 | -0.6144497 | 71 | ACTG1 |
| VCL | 26 | -0.3727195 | -0.6131192 | 7414 | VCL |
| FAM120A | 27 | -0.3285104 | -0.6090931 | 23196 | FAM120A |

| Gene | Rank | DMSO | Treat | EntrezID | Symbol |
| --- | --- | --- | --- | --- | --- |
| HSP90AB1 | 28 | -0.3253732 | -0.605722 | 3326 | HSP90AB1 |
| GTF2H5 | 29 | -0.1308977 | -0.6055634 | 404672 | GTF2H5 |
| APOLD1 | 30 | -0.3387373 | -0.6045638 | 81575 | APOLD1 |
| PPP1R15B | 31 | -0.2944295 | -0.6036814 | 84919 | PPP1R15B |
| FAM133B | 32 | -0.1080237 | -0.6026955 | 257415 | FAM133B |
| UVSSA | 33 | -0.2945939 | -0.6026266 | 57654 | UVSSA |
| NBN | 34 | -0.2980664 | -0.602537 | 4683 | NBN |
| RAD18 | 35 | -0.0184755 | -0.5978491 | 56852 | RAD18 |
| ERCC8 | 36 | -0.223341 | -0.5904588 | 1161 | ERCC8 |
| ETAA1 | 37 | -0.2938704 | -0.5789183 | 54465 | ETAA1 |
| SPRTN | 38 | -0.3473528 | -0.5750508 | 83932 | SPRTN |
| FLII | 39 | -0.3271424 | -0.5702527 | 2314 | FLII |
| SRP54 | 40 | -0.3080039 | -0.5686671 | 6729 | SRP54 |
| GSS | 41 | -0.2006774 | -0.5637586 | 2937 | GSS |
| SLC44A3 | 42 | -0.1966919 | -0.5589121 | 126969 | SLC44A3 |
| GTF3C1 | 43 | -0.2859651 | -0.5576919 | 2975 | GTF3C1 |
| BARD1 | 44 | -0.2596777 | -0.5570577 | 580 | BARD1 |
| UBE2K | 45 | -0.1413154 | -0.5570508 | 3093 | UBE2K |
| SLC7A1 | 46 | -0.3216968 | -0.555865 | 6541 | SLC7A1 |
| SUCO | 47 | -0.2802828 | -0.5547689 | 51430 | SUCO |
| RUNX2 | 48 | -0.0624873 | -0.5533901 | 860 | RUNX2 |
| SECISBP2 | 49 | -0.3153765 | -0.5495571 | 79048 | SECISBP2 |
| RNF113A | 50 | -0.3141335 | -0.5487987 | 7737 | RNF113A |
| PSMF1 | 51 | -0.3008089 | -0.5485437 | 9491 | PSMF1 |
| CNPY2 | 52 | -0.3117067 | -0.5441522 | 10330 | CNPY2 |
| SCAF4 | 53 | -0.2546991 | -0.54176 | 57466 | SCAF4 |
| DUSP10 | 54 | -0.2936929 | -0.5357761 | 11221 | DUSP10 |
| CCS | 55 | -0.1684249 | -0.5313915 | 9973 | CCS |
| OTUB1 | 56 | -0.2843012 | -0.5311571 | 55611 | OTUB1 |
| MCM9 | 57 | -0.1977179 | -0.5297439 | 254394 | MCM9 |
| POLR2A | 58 | -0.0658533 | -0.52467 | 5430 | POLR2A |
| TMEM53 | 59 | -0.2582045 | -0.519789 | 79639 | TMEM53 |
| HACE1 | 60 | -0.2943966 | -0.5181 | 57531 | HACE1 |
| RXFP4 | 61 | -0.2558961 | -0.5165972 | 339403 | RXFP4 |
| EME1 | 62 | -0.215416 | -0.5140326 | 146956 | EME1 |
| BCL2L1 | 63 | -0.1415258 | -0.509979 | 598 | BCL2L1 |
| CDC25A | 64 | -0.1635515 | -0.5080418 | 993 | CDC25A |
| UPF1 | 65 | -0.2318316 | -0.5075868 | 5976 | UPF1 |
| HSPB3 | 66 | -0.2446037 | -0.507311 | 8988 | HSPB3 |
| MTERFD2 | 67 | -0.2726669 | -0.5065803 | 130916 | MTERFD2 |
| UPF3A | 68 | -0.1999803 | -0.5050429 | 65110 | UPF3A |
| FAM175A | 69 | -0.2782374 | -0.5034297 | 84142 | FAM175A |
| ADCY8 | 70 | -0.2545478 | -0.4966254 | 114 | ADCY8 |
| ZFAND3 | 71 | -0.214094 | -0.4924959 | 60685 | ZFAND3 |
| DNAJA1 | 72 | -0.2530878 | -0.4921719 | 3301 | DNAJA1 |
| XPC | 73 | -0.1525156 | -0.4877805 | 7508 | XPC |
| RUNX1 | 74 | -0.1905557 | -0.4869877 | 861 | RUNX1 |
| CYB5R4 | 75 | -0.2497534 | -0.4851677 | 51167 | CYB5R4 |
| GPA33 | 76 | -0.2143834 | -0.4832167 | 10223 | GPA33 |
| ADAMTSL4 | 77 | -0.2383558 | -0.4788046 | 54507 | ADAMTSL4 |
| PPIA | 78 | -0.1914239 | -0.4696081 | 5478 | PPIA |
| CHD2 | 79 | -0.2106807 | -0.4691807 | 1106 | CHD2 |
| MKS1 | 80 | -0.1291878 | -0.4590328 | 54903 | MKS1 |
| ROPN1L | 81 | -0.1204538 | -0.4581986 | 83853 | ROPN1L |
| DUSP28 | 82 | -0.0917856 | -0.4569094 | 285193 | DUSP28 |
| DNMT1 | 83 | -0.1663992 | -0.4556341 | 1786 | DNMT1 |
| MNT | 84 | -0.0885038 | -0.4551584 | 4335 | MNT |
| PAN2 | 85 | -0.187149 | -0.4540898 | 9924 | PAN2 |
| SERP1 | 86 | -0.2148306 | -0.4492985 | 27230 | SERP1 |
| ACER2 | 87 | -0.1507793 | -0.4476095 | 340485 | ACER2 |
| HIST1H3B | 88 | -0.2083854 | -0.4475957 | 8358 | HIST1H3B |
| C1orf52 | 89 | -0.1876751 | -0.4466513 | 148423 | C1orf52 |
| SMARCC1 | 90 | -0.0559303 | -0.4463617 | 6599 | SMARCC1 |
| MUSTN1 | 91 | -0.1604998 | -0.4447141 | 389125 | MUSTN1 |
| C1orf112 | 92 | -0.0500467 | -0.4446658 | 55732 | C1orf112 |
| PKN3 | 93 | -0.0802105 | -0.4379304 | 29941 | PKN3 |
| UIMC1 | 94 | -0.1941138 | -0.4334976 | 51720 | UIMC1 |
| FAM178A | 95 | -0.1695429 | -0.4299542 | 55719 | FAM178A |
| FBXO7 | 96 | -0.1673397 | -0.4282031 | 25793 | FBXO7 |

**Table 4.** CRISPR essential gene list for Doxorubicin treatment.

| **Gene** | **Rank** | **DMSO** | **Treat** | **EntrezID** | **Symbol** |
| --- | --- | --- | --- | --- | --- |
| ABCC1 | 1 | -0.0742760 | -1.5151284 | 4363 | ABCC1 |
| TDP2 | 2 | -0.3092421 | -1.1145884 | 51567 | TDP2 |
| GCLM | 3 | -0.3146230 | -0.8633248 | 2730 | GCLM |
| ZNF451 | 4 | 0.0425952 | -0.8365990 | 26036 | ZNF451 |
| TAS2R30 | 5 | -0.1359444 | -0.7913176 | 259293 | TAS2R30 |
| UIMC1 | 6 | -0.2276415 | -0.7744236 | 51720 | UIMC1 |
| FAM175A | 7 | -0.2786786 | -0.7236031 | 84142 | FAM175A |
| ATM | 8 | -0.3411623 | -0.6845115 | 472 | ATM |
| HNRNPR | 9 | -0.3712627 | -0.6511459 | 10236 | HNRNPR |
| HIST1H2BJ | 10 | -0.2855531 | -0.6429758 | 8970 | HIST1H2BJ |
| BABAM1 | 11 | -0.2803874 | -0.6309492 | 29086 | BABAM1 |
| DYNLL1 | 12 | -0.2048135 | -0.6206121 | 8655 | DYNLL1 |
| NHEJ1 | 13 | -0.3072919 | -0.6050336 | 79840 | NHEJ1 |
| OSGEP | 14 | -0.3534633 | -0.5985460 | 55644 | OSGEP |
| POLR3D | 15 | -0.3618902 | -0.5802603 | 661 | POLR3D |
| SUMO2 | 16 | -0.3706170 | -0.5712525 | 6613 | SUMO2 |
| ZEB2 | 17 | -0.3295330 | -0.5706086 | 9839 | ZEB2 |
| SPTLC2 | 18 | -0.3430146 | -0.5687876 | 9517 | SPTLC2 |
| PPP2R5D | 19 | -0.3761414 | -0.5667521 | 5528 | PPP2R5D |
| POLQ | 20 | -0.3550222 | -0.5592744 | 10721 | POLQ |
| FANCL | 21 | -0.3718562 | -0.5590113 | 55120 | FANCL |
| BRE | 22 | -0.2885273 | -0.5556048 | 9577 | BRE |
| CRAMP1L | 23 | 0.1005674 | -0.5450668 | 57585 | CRAMP1L |
| PSMF1 | 24 | -0.3135403 | -0.5438067 | 9491 | PSMF1 |
| ANKRD35 | 25 | -0.3452322 | -0.5349789 | 148741 | ANKRD35 |
| SNAPC1 | 26 | -0.3400404 | -0.5343073 | 6617 | SNAPC1 |
| POLR2J | 27 | -0.3702257 | -0.5341826 | 5439 | POLR2J |

| Gene | Rank | DMSO | Treat | EntrezID | Symbol |
| --- | --- | --- | --- | --- | --- |
| DENR | 28 | -0.3627315 | -0.5307000 | 8562 | DENR |
| CDKN2AIP | 29 | -0.3488456 | -0.5294883 | 55602 | CDKN2AIP |
| BRCC3 | 30 | -0.1904970 | -0.5288721 | 79184 | BRCC3 |
| ZMAT2 | 31 | -0.3605987 | -0.5260403 | 153527 | ZMAT2 |
| CDC25B | 32 | -0.3444430 | -0.5220522 | 994 | CDC25B |
| FBLIM1 | 33 | -0.3246935 | -0.5216645 | 54751 | FBLIM1 |
| TMEM230 | 34 | -0.3671732 | -0.5203005 | 29058 | TMEM230 |
| HNRNPH1 | 35 | -0.3185821 | -0.5189365 | 3187 | HNRNPH1 |
| SFR1 | 36 | -0.1113358 | -0.5187288 | 119392 | SFR1 |
| XRCC1 | 37 | -0.3541678 | -0.5175725 | 7515 | XRCC1 |
| ACTG1 | 38 | -0.3619945 | -0.5153015 | 71 | ACTG1 |
| ATP13A1 | 39 | -0.3335899 | -0.5134113 | 57130 | ATP13A1 |
| WDR24 | 40 | -0.3290373 | -0.5119574 | 84219 | WDR24 |
| OPRD1 | 41 | -0.3228020 | -0.5072284 | 4985 | OPRD1 |
| C10orf76 | 42 | -0.2693973 | -0.5047012 | 79591 | C10orf76 |
| IRS1 | 43 | -0.1707344 | -0.5039396 | 3667 | IRS1 |
| TRIT1 | 44 | -0.3028437 | -0.5015371 | 54802 | TRIT1 |
| SECISBP2 | 45 | -0.3226454 | -0.4992730 | 79048 | SECISBP2 |
| OR5K2 | 46 | -0.2687973 | -0.4988507 | 402135 | OR5K2 |
| ETAA1 | 47 | -0.3146426 | -0.4977775 | 54465 | ETAA1 |
| FAF2 | 48 | -0.3267154 | -0.4976390 | 23197 | FAF2 |
| RAB14 | 49 | -0.3079703 | -0.4974105 | 51552 | RAB14 |
| GSS | 50 | -0.2274002 | -0.4970020 | 2937 | GSS |
| RHOT2 | 51 | -0.3358727 | -0.4920861 | 89941 | RHOT2 |
| TP53BP1 | 52 | -0.2810918 | -0.4902860 | 7158 | TP53BP1 |
| SCAF4 | 53 | -0.2746217 | -0.4841169 | 57466 | SCAF4 |
| NBN | 54 | -0.3041417 | -0.4821990 | 4683 | NBN |
| ANKRD17 | 55 | -0.3014153 | -0.4800042 | 26057 | ANKRD17 |
| NFE2L2 | 56 | -0.2954866 | -0.4797411 | 4780 | NFE2L2 |
| AP1B1 | 57 | -0.2068745 | -0.4753791 | 162 | AP1B1 |
| SKP2 | 58 | -0.2945930 | -0.4744998 | 6502 | SKP2 |
| ATP5D | 59 | -0.3115119 | -0.4717718 | 513 | ATP5D |
| MRPL24 | 60 | -0.2615575 | -0.4714464 | 79590 | MRPL24 |
| APOL4 | 61 | -0.2976585 | -0.4713148 | 80832 | APOL4 |
| AIP | 62 | -0.3046113 | -0.4688430 | 9049 | AIP |
| NFYA | 63 | -0.3156535 | -0.4687946 | 4800 | NFYA |
| AKT1S1 | 64 | -0.2680733 | -0.4663505 | 84335 | AKT1S1 |
| UBE2A | 65 | -0.3044743 | -0.4652358 | 7319 | UBE2A |
| C17orf85 | 66 | -0.3022698 | -0.4640726 | 55421 | C17orf85 |
| FAM69A | 67 | -0.2966084 | -0.4625424 | 388650 | FAM69A |
| SERP1 | 68 | -0.2180994 | -0.4623970 | 27230 | SERP1 |
| PABPC1 | 69 | -0.1802178 | -0.4594059 | 26986 | PABPC1 |
| NDUFB9 | 70 | -0.1906992 | -0.4572250 | 4715 | NDUFB9 |
| MORC2 | 71 | -0.2659666 | -0.4563733 | 22880 | MORC2 |
| DPY30 | 72 | -0.2926885 | -0.4551824 | 84661 | DPY30 |
| ESPN | 73 | -0.2385860 | -0.4530499 | 83715 | ESPN |
| SUCO | 74 | -0.2775959 | -0.4496434 | 51430 | SUCO |
| PCDH7 | 75 | -0.2863749 | -0.4489649 | 5099 | PCDH7 |
| USP24 | 76 | -0.2407188 | -0.4424704 | 23358 | USP24 |
| CCDC47 | 77 | -0.1620402 | -0.4409818 | 57003 | CCDC47 |
| CDK5 | 78 | -0.1117336 | -0.4394170 | 1020 | CDK5 |
| CHD8 | 79 | -0.2042917 | -0.4377138 | 57680 | CHD8 |
| KRTAP5-11 | 80 | -0.1198474 | -0.4373260 | 440051 | KRTAP5-11 |
| SLC44A3 | 81 | -0.1998304 | -0.4330541 | 126969 | SLC44A3 |
| MED10 | 82 | -0.2738977 | -0.4314824 | 84246 | MED10 |
| CDCA3 | 83 | -0.2657057 | -0.4310600 | 83461 | CDCA3 |
| TOMM70A | 84 | -0.2406601 | -0.4256595 | 9868 | TOMM70A |
| SNAPIN | 85 | -0.2670167 | -0.4244617 | 23557 | SNAPIN |
| C17orf58 | 86 | -0.2170297 | -0.4236031 | 284018 | C17orf58 |
| PRKCG | 87 | -0.1466280 | -0.4232846 | 5582 | PRKCG |
| NDUFA11 | 88 | -0.2080877 | -0.4187773 | 126328 | NDUFA11 |
| RBM28 | 89 | -0.2656405 | -0.4184172 | 55131 | RBM28 |
| FKBP3 | 90 | -0.2503913 | -0.4184103 | 2287 | FKBP3 |
| C2CD4D | 91 | -0.1885860 | -0.4157031 | 100191040 | C2CD4D |
| DNM1 | 92 | -0.0784177 | -0.4098318 | 1759 | DNM1 |
| IQCH | 93 | -0.2148056 | -0.4092502 | 64799 | IQCH |

### Deficiency of essential genes contribute to increased chemosensitivity in TNBC

In our study, a cell survival assay using siRNA-mediated gene silencing was performed to validate the novel genes identified from our CRISPR essential gene list. The results, as illustrated in the Figure 6A-6C, demonstrate that knocking down the expression of specific genes, namely ATR, NEPPS, and MCM9, increases the sensitivity of triple-negative breast cancer (TNBC) cell lines to the chemotherapeutic drug cisplatin.

**Figure 5.**
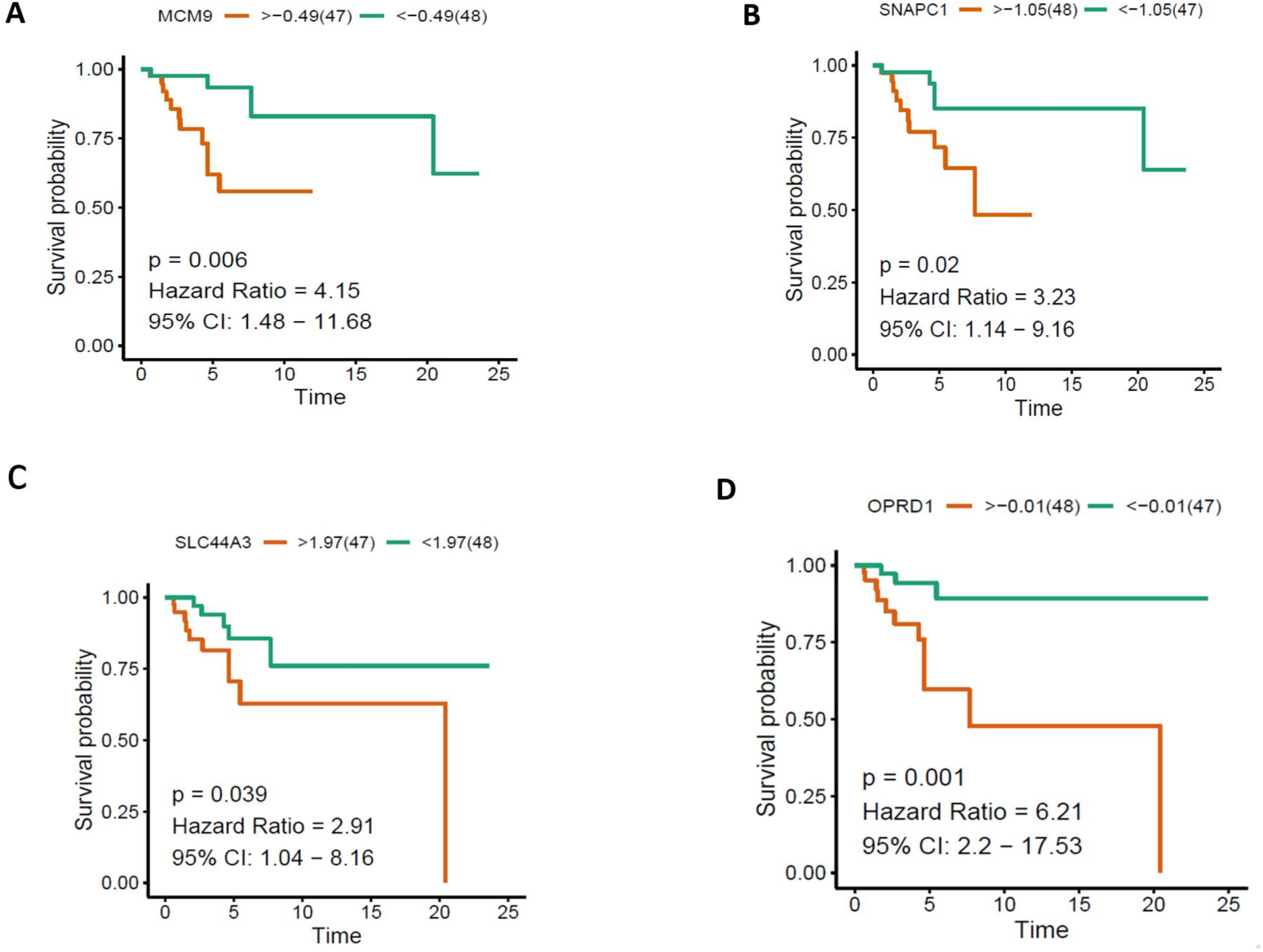
Kaplan-Meier overall survival analysis of essential genes based on TCGA database. Overall survival analysis for essential genes MCM9, SNAPC1,SLC44A3,OPRD1 (A-D) TNBC.

**Figure 6.**
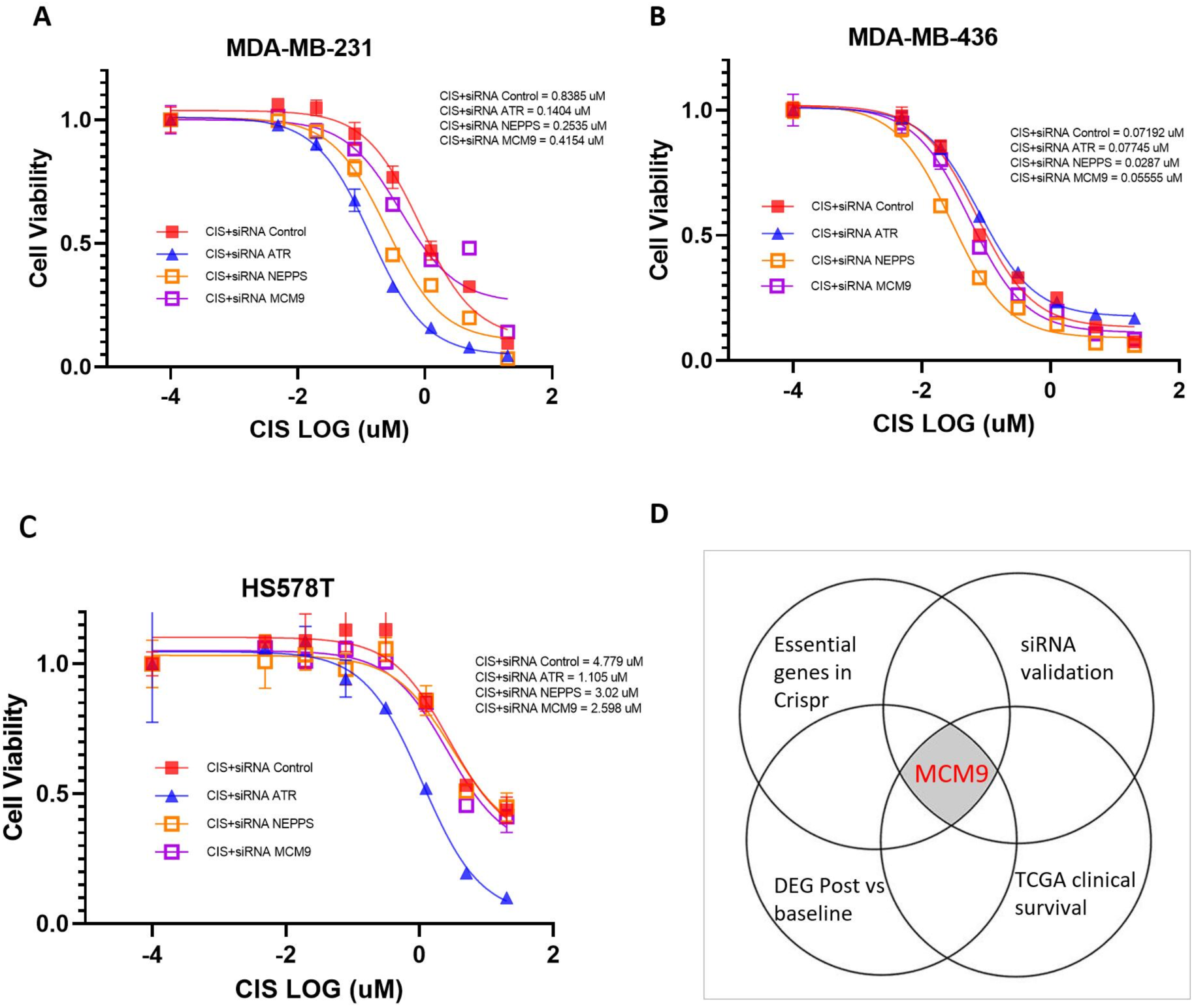
siRNA Knockdown cell viability assay. Cell growth inhibition of MDA-MB-231, MDA-MB-436 and HS578T transfected with siRNA ATR, siRNA NEPPS or siRNA MCM9, followed by treatment with 4-serial-diluted cisplatin doses for 120 h (A-C). A Venn diagram can be created by overlapping different essential gene validation methods (D).

### Relationship between essential gene expression and TNBC patient survival outcomes

In the essential gene lists identified through our CRISPR screening, we performed an overall survival analysis using TCGA data for those genes (Figure 5)that were not well-studied previously. The TCGA data unveiled notable differences in overall survival between patients exhibiting high and low expression levels of the following genes (Figure 3D-E): MCM9 (Hazard Ratio = 4.15, 95% CI: 1.48-11.68, p-value = 0.006), SNAPC1 (Hazard Ratio = 3.23, 95% CI: 1.14-9.16, p-value = 0.02), SLC44A3 (Hazard Ratio = 2.91, 95% CI: 1.04-8.16, p-value = 0.039), and OPRD1 (Hazard Ratio = 6.21, 95% CI: 2.2-17.53, p-value = 0.001). The Kaplan-Meier curves showed a distinct separation between the high and low expression groups for each gene, indicating that gene expression levels are correlated with patient outcomes in the cancer type under investigation.

## DISCUSSION

Upon analyzing the data from our CRISPR screening, we found that both the read level QC and sample level QC demonstrated high data quality. Missed sgRNAs are not detected in the final sequencing data after performing a CRISPR-Cas9 screen. This can occur for various reasons, such as the low initial representation of the sgRNA in the library, poor experimental conditions, or technical issues during sequencing ^30^. For all our samples, the number of missed sgRNAs is below 500, a tiny proportion compared to the 70,948 sgRNAs in the tkov3 library. This indicates that our experimental steps were successful and without significant issues. The Gini index, also known as the Gini coefficient, is a statistical measure that quantifies the inequality or dispersion of a distribution. In the context of CRISPR-Cas9 screens, the Gini index assesses the unevenness of sgRNA representation in the initial library or after selection ^31^. The overall Gini index for all samples is low, consistent with the expected Gini index in negative screening. Furthermore, the normalization results of the three datasets are consistent, indicating that our results are comparable across samples, allowing for more accurate identification of essential genes.

PCA (Principal Component Analysis) plots are valuable for visualizing high-dimensional data in a lower-dimensional space—the first three principal components in our analysis account for over 80% of the sample variance. Through the 3D PCA plot, a clear pattern can be observed, with triplicate samples from each group clustering together. The pairwise sample correlation results show that the correlation value within each group is greater than 0.9, while the correlation value between groups does not exceed 0.82. This demonstrates good reproducibility in our experiments, and the data is trustworthy and also There are significant differences between groups under different experimental conditions.

In our study, we used genome-wide CRISPR screening to identify essential genes that could help overcome cisplatin/doxorubicin resistance in triple-negative breast cancer. Our results were supported by a gene table (Table 1 for CIS and Table 2 for DOX treatment group) including summary of well-studied genes from previous research.

For cisplatin treatment, the well-studied genes included in our gene list are DNMT1, PPIA, RUNX, BCL2L1, RUNX2, NBN, GTF2H5, USP22, HSP90AB1, CDC25B, NCF1, FANCA, FANCG, and ERCC1. Some of these genes have known drug targets or inhibitors. For example, DNMT1, involved in the DNA methylation pathway, is targeted by decitabine, which is clinically used for other cancer types ^32^. RUNX, which regulates apoptosis and cell proliferation, has a small molecule inhibitor (AI-10-49) in preclinical development ^33^. BCL2L1, involved in the intrinsic apoptotic pathway, can be targeted by BikDD and lapatinib, both in preclinical stages ^32^. RUNX2, which also plays a role in regulating the cell cycle and apoptosis, is targeted by BET inhibitors JQ1 and I-BET762, currently in phase I/II clinical trials ^34^.

Furthermore, several genes in our list have been previously implicated in chemoresistance. For example, PPIA is involved in miRNA regulation and impacts breast cancer cell sensitivity to doxorubicin ^35^. RUNX is involved in the YAP signaling pathway, and its knockdown enhances sensitivity to doxorubicin in breast cancer cells ^32^. NBN is involved in DNA repair and homologous recombination, playing a role in doxorubicin, paclitaxel, and carboplatin resistance in HER2- and MDM2-enriched breast cancer subtypes ^36^. GTF2H5 is involved in nucleotide.

For doxorubicin treatment, the well-studied genes included in our gene list are ABCC1, HIST1H2BJ, ZEB2, ATM, FANCL, CDC25B, XRCC1, ACTG1, IRS1, NBN, NFE2L2, NDUFB9, CDK5, and CDCA3. Several genes in this list have previously been reported to play a role in chemoresistance. For instance, ABCC1 is a drug efflux transporter that has been implicated in resistance to doxorubicin, paclitaxel, and cisplatin in TNBC ^37^. HIST1H2BJ is involved in glutathione synthesis and copper chelation, promoting resistance to doxorubicin, paclitaxel, and cisplatin in TNBC ^38^. ZEB2, a transcription factor, is associated with drug resistance in breast cancer cells by regulating the epithelial-mesenchymal transition (EMT) ^39^. ATM, a kinase involved in DNA damage response, is known to contribute to doxorubicin resistance in breast cancer cells.FANCL is part of the Fanconi anemia DNA repair pathway, and its inhibition has been shown to enhance the sensitivity of breast cancer cells to cisplatin and olaparib. CDC25B, a cell cycle regulator, has been implicated in doxorubicin and paclitaxel resistance in breast cancer cells ^40^. XRCC1, a key protein in base excision repair, has been reported to contribute to resistance against doxorubicin in breast cancer cells.

Some of these genes have potential drug targets or inhibitors. The ATM kinase inhibitor KU-55933, which targets ATM involved in the DNA damage response pathway, is in preclinical development ^41^. FANCL, part of the Fanconi anemia DNA repair pathway, has been targeted by small molecule inhibitors such as curcumin in preclinical studies ^42^. CDK5 inhibitors, like roscovitine and dinaciclib, have shown promise in preclinical studies and are in clinical trials for various cancer types ^43^. CDK5 is involved in cell cycle regulation and the DNA damage response, which contribute to chemoresistance.

CDC25B and NBN are the overlapping genes between the cisplatin and doxorubicin essential gene lists. CDC25B is involved in cell cycle regulation and DNA damage response and has been targeted by Thiostrepton, FDI-6, and Siomycin A in preclinical studies for platinum-resistant ovarian cancer treatment ^44^. NBN, on the other hand, is involved in DNA repair and homologous recombination and has been studied in vitro using siRNA. While no drug targets or inhibitors have been identified for NBN, its role in DNA repair suggests it may be a potential therapeutic target in the future.

In our study, the KEGG enrichment analysis of the essential gene list revealed distinct, yet overlapping, pathways enriched in the cisplatin and doxorubicin treatment groups. Both groups showed a strong connection to DNA damage repair, highlighting the importance of these pathways in response to chemotherapeutic agents. In the cisplatin treatment group, the top 15 enriched pathways were primarily associated with double-strand break repair, response to radiation, DNA replication, regulation of DNA metabolic process, and nucleotide-excision repair. These results suggest that cisplatin-induced DNA damage triggers a range of cellular responses, including the activation of DNA repair mechanisms and changes in DNA conformation, which may contribute to chemoresistance.

In contrast, the doxorubicin treatment group’s top 15 enriched pathways were mainly involved in double-strand break repair via nonhomologous end joining, non-recombinational repair, regulation of response to DNA damage stimulus, cell cycle G2/M phase transition, and positive regulation of DNA metabolic process. These findings indicate that doxorubicin-induced DNA damage elicits different cellular responses, primarily focusing on cell cycle regulation and checkpoint activation. This disparity in pathway enrichment between the two treatment groups underscores the differences in the mechanisms of action of cisplatin and doxorubicin and the cellular responses they induce.

Our study has provided valuable insights into the distinct yet overlapping pathways involved in the cellular response to cisplatin and doxorubicin treatment. These results emphasize the importance of understanding the molecular mechanisms underlying DNA damage repair and response to chemotherapeutic agents to develop more effective therapies and overcome chemoresistance in cancer treatment. Further studies are warranted to investigate the potential therapeutic targets identified in these enriched pathways and to explore the crosstalk between them to develop novel strategies for overcoming chemoresistance in cancer cells.

Our cell survival assay demonstrates that knocking down the expression of specific genes, namely ATR, NEPPS, and MCM9, would increase the sensitivity of triple-negative breast cancer (TNBC) cell lines to the chemotherapeutic drug cisplatin. We employed not only the MDA-MB-231 cell line, which was used for our initial screening, but also other TNBC cell lines, such as MDA-MB-436 and HS578T, to strengthen our hypothesis. Our results showed a significant reduction in cell viability in all three cell lines following the knockdown of ATR, NEPPS, and MCM9, indicating an increased sensitivity to cisplatin treatment.

In our survival analysis, we observed a significant association between the expression levels of MCM9, SNAPC1, SLC44A3, and OPRD1 genes and overall survival in the patient population analyzed using TCGA data. Specifically, patients with lower expression levels of these genes experienced better survival outcomes, suggesting a potential therapeutic strategy that involves downregulating these genes to enhance patients’ response to chemotherapy. These findings are consistent with the results of our CRISPR screening. A Venn diagram (Figure 5H) can be created by overlapping different essential gene validation methods, such as genome wide CRISPR screening, siRNA knockdown validation, differential gene expression analysis, and survival analysis. The intersection of these methods helps identify the core set of essential genes, such as MCM9, with a higher level of confidence as a potential gene target for overcoming triple-negative breast cancer.

In conclusion, our study strongly supports the notion that utilizing CRISPR screening for essential cancer genes is an efficient and practical approach to identify potential drug targets to overcome chemotherapeutic drug resistance. By integrating multiple data analysis techniques and biological experimental analyses, we have identified several genes that hold promise as potential drug targets for combating chemoresistance. Our findings underscore the importance of leveraging advanced genetic screening tools and data-driven methods to better understand the molecular mechanisms underlying drug resistance and to develop more effective therapeutic strategies for cancer treatment.

## MATERIALS AND METHODS

### Cell Culture

Human triple-negative breast cancer cell lines MDA-MB-231, MDA-MB-436, and HS-578T, along with the Homo sapiens embryonic kidney cell line 293T, were obtained from the American Type Culture Collection (Manassas, VA, USA) for this study. All cell lines were cultured in Ham’s F-12K (Kaighn’s) medium, supplemented with 10% fetal bovine serum (VWR, Radnor, PA, USA), 1% GlutMax, 1% sodium pyruvate, and penicillin-streptomycin (Gibco, Waltham, MA, USA). The cell lines were incubated at 37 °C in a 5% CO2 atmosphere. All cell lines underwent authentication via STR profiling and were tested for mycoplasma contamination every three months.

### TKOv3 library Construction

The Toronto Knock-Out CRISPR (TKOv3) library, containing 71,090 sgRNAs targeting 18,049 protein-coding genes, was acquired from Addgene (Watertown, MA, USA) and expanded 1000-fold using the electroporation method. For lentivirus production, 7.5×10^6 293T cells were seeded in 15 cm plates and prepared for transfection. Following the manufacturer’s guidelines, packaging vectors psPAX2, pMD2.G (Addgene), TKOv3 library plasmid, and Lipofectamine (Thermo Fisher Scientific) were mixed in OptiMEM (Thermo Fisher Scientific, Waltham, MA, USA). After 48 h of incubation, the lentivirus-containing medium was collected and stored at −80°C.

### Pooled sgRNA screens

MDA-MB-231 cells were transduced with the TKOv3 lentivirus library at a low multiplicity of infection (MOI) of 0.3. After 72 h of puromycin (2 μg/mL) selection, surviving cells were considered baseline samples (T0), and 3×10^7 cells were harvested and stored at −80°C. The remaining cells were divided into three groups (control, cisplatin treatment, and doxorubicin treatment), each performed in triplicate. Following four weeks of treatment, 3×10^7 cells were harvested from each group. Genomic DNA was extracted using the QIAamp Blood Maxi Kit (Qiagen, Hilden, Germany). Two polymerase chain reactions (PCRs) were carried out to enrich the sgRNA-targeted genomic regions and amplify the sgRNA. The resulting libraries were sequenced on a NovaSeq 6000 system (Illumina, San Diego, CA, USA), producing nearly 80 million reads per sample to achieve 600x coverage of the CRISPR library.

### Crispr Data analysis

CRISPR pooled CRISPR/Cas9 knockout screening data was analyzed using the MAGeCK (Model-based Analysis of Genome-wide CRISPR-Cas9 Knockout) algorithm. Data were first normalized using a list of non-targeting control sgRNAs. Gene essentiality scores (beta-scores) were then determined for each group using the MAGeCK MLE (Maximum Likelihood Estimation) method. Principal component analysis (PCA) and Pearson correlation analysis were performed using the R packages “stats” and “corrplot”, respectively.

### Cell survival assay using siRNA-mediated gene silencing

Small interfering RNAs (siRNAs) were used to validate the essential genes identified in our study. These siRNAs, which were specifically designed to target the essential genes, were purchased from Thermo Fisher Scientific (MA, USA). Detailed information on the siRNAs is provided in the Supplementary SI Gene list. We transfected MDA-MB-231, MDA-MB-436, and HS578T cells with siRNAs using the Lipofectamine RNAiMAX Transfection Reagent kit (#13778150, Thermo Fisher Scientific, MA, USA) according to the manufacturer’s protocol. The cells were then seeded into 96-well plates at a density of 2.5 x 10^3 cells per well. After 24 hours, the medium was replaced, and the cells were treated with cisplatin and doxorubicin. Following 120 hours of incubation, we assessed cell viability using the alamarBlue HS Cell Viability Reagent (#A50100, Thermo Fisher Scientific, MA, USA). The absorbance of each well was measured using a microplate reader. We determined the half-maximal inhibitory concentration (IC50) values with the help of GraphPad Prism 7 software.

### 3.5.6 Survival analysis

In this study, we utilized The Cancer Genome Atlas (TCGA) database to analyze the overall survival of TNBC patients with varying gene expression levels. The gene expression values were obtained from the TCGA database, and patients were divided into two groups based on the median expression value: high expression group and low expression group. Kaplan-Meier curves were generated to compare the overall survival of the two groups. Hazard ratios and p-values were calculated using the log-rank test to assess the statistical significance of the differences in survival.

## Reference

1 Ades, F. et al. Luminal B breast cancer: molecular characterization, clinical management, and future perspectives. Journal of clinical oncology 32, 2794–2803 (2014).

2 Shao, S. et al. Site-specific and hydrophilic ADCs through disulfide-bridged linker and branched PEG. Bioorganic & Medicinal Chemistry Letters 28, 1363–1370 (2018).

3 Nedeljković, M. & Damjanović, A. Mechanisms of chemotherapy resistance in triple-negative breast cancer—how we can rise to the challenge. Cells 8, 957 (2019).

4 Yagata, H., Kajiura, Y. & Yamauchi, H. Current strategy for triple-negative breast cancer: appropriate combination of surgery, radiation, and chemotherapy. Breast Cancer 18, 165–173 (2011).

5 Dasari, S. & Tchounwou, P. B. Cisplatin in cancer therapy: molecular mechanisms of action. European journal of pharmacology 740, 364–378 (2014).

6 Qi, R. et al. Sequence Dependent Repair of 1, N 6-Ethenoadenine by DNA Repair Enzymes ALKBH2, ALKBH3, and AlkB. Molecules 26, 5285 (2021).

7 Rivankar, S. An overview of doxorubicin formulations in cancer therapy. Journal of cancer research and therapeutics 10, 853–858 (2014).

8 Keam, B. et al. Prognostic impact of clinicopathologic parameters in stage II/III breast cancer treated with neoadjuvant docetaxel and doxorubicin chemotherapy: paradoxical features of the triple negative breast cancer. BMC cancer 7, 1–11 (2007).

9 Byrski, T. et al. Response to neoadjuvant therapy with cisplatin in BRCA1-positive breast cancer patients. Breast cancer research and treatment 115, 359–363 (2009).

10 Sikov, W. M. et al. Impact of the addition of carboplatin and/or bevacizumab to neoadjuvant once-per-week paclitaxel followed by dose-dense doxorubicin and cyclophosphamide on pathologic complete response rates in stage II to III triple-negative breast cancer: CALGB 40603 (Alliance). Journal of clinical oncology 33, 13 (2015).

11 Loibl, S. et al. Addition of the PARP inhibitor veliparib plus carboplatin or carboplatin alone to standard neoadjuvant chemotherapy in triple-negative breast cancer (BrighTNess): a randomised, phase 3 trial. The Lancet Oncology 19, 497–509 (2018).

12 Schmid, P. et al. Pembrolizumab for early triple-negative breast cancer. New England Journal of Medicine 382, 810–821 (2020).

13 Schmid, P. et al. Atezolizumab and nab-paclitaxel in advanced triple-negative breast cancer. New England Journal of Medicine 379, 2108–2121 (2018).

14 Dent, R. A. et al. Phase I trial of the oral PARP inhibitor olaparib in combination with paclitaxel for first-or second-line treatment of patients with metastatic triple-negative breast cancer. Breast cancer research 15, 1–8 (2013).

15 Geenen, J. J., Linn, S. C., Beijnen, J. H. & Schellens, J. H. PARP inhibitors in the treatment of triple-negative breast cancer. Clinical pharmacokinetics 57, 427–437 (2018).

16 Nagayama, A., Vidula, N., Ellisen, L. & Bardia, A. Novel antibody–drug conjugates for triple negative breast cancer. Therapeutic advances in medical oncology 12, 1758835920915980 (2020).

17 Lyons, T. G. Targeted therapies for triple-negative breast cancer. Current treatment options in oncology 20, 1–13 (2019).

18 Bradbury, A., Hall, S., Curtin, N. & Drew, Y. Targeting ATR as Cancer Therapy: A new era for synthetic lethality and synergistic combinations? Pharmacology & therapeutics 207, 107450 (2020).

19 Cruz, C. et al. RAD51 foci as a functional biomarker of homologous recombination repair and PARP inhibitor resistance in germline BRCA-mutated breast cancer. Annals of Oncology 29, 1203–1210 (2018).

20 Cortesi, L., Rugo, H. S. & Jackisch, C. An overview of PARP inhibitors for the treatment of breast cancer. Targeted oncology 16, 255–282 (2021).

21 Ström, C. E. et al. Poly (ADP-ribose) polymerase (PARP) is not involved in base excision repair but PARP inhibition traps a single-strand intermediate. Nucleic acids research 39, 3166–3175 (2011).

22 Yoshida, K. & Miki, Y. Role of BRCA1 and BRCA2 as regulators of DNA repair, transcription, and cell cycle in response to DNA damage. Cancer science 95, 866–871 (2004).

23 Lord, C. J. & Ashworth, A. PARP inhibitors: Synthetic lethality in the clinic. Science 355, 1152–1158 (2017).

24 Ran, F. A. et al. Genome engineering using the CRISPR-Cas9 system. Nature protocols 8, 2281–2308 (2013).

25 Tang, S. et al. Generation of dual-gRNA library for combinatorial CRISPR screening of synthetic lethal gene pairs. STAR protocols 3, 101556 (2022).

26. Li, C., Fu, J., Shao, S. & Luo, Z.-Q. Legionella pneumophila exploits the endo-lysosomal network for phagosome biogenesis by co-opting SUMOylated Rab7. bioRxiv (2023).

27 Tang, S. et al. Synthetic lethal gene pairs: Experimental approaches and predictive models. Frontiers in Genetics 13, 961611 (2022).

28 Sanjana, N. E., Shalem, O. & Zhang, F. Improved vectors and genome-wide libraries for CRISPR screening. Nature methods 11, 783–784 (2014).

29 Dai, M. et al. In vivo genome-wide CRISPR screen reveals breast cancer vulnerabilities and synergistic mTOR/Hippo targeted combination therapy. Nature Communications 12, 3055 (2021).

30 Wang, B. et al. Integrative analysis of pooled CRISPR genetic screens using MAGeCKFlute. Nature protocols 14, 756–780 (2019).

31 Hart, T., Brown, K. R., Sircoulomb, F., Rottapel, R. & Moffat, J. Measuring error rates in genomic perturbation screens: gold standards for human functional genomics. Molecular systems biology 10, 733 (2014).

32 Sharma, S., Kelly, T. K. & Jones, P. A. Epigenetics in cancer. Carcinogenesis 31, 27–36 (2010).

33 Samarakkody, A. S., Shin, N.-Y. & Cantor, A. B. Role of RUNX family transcription factors in DNA damage response. Molecules and cells 43, 99 (2020).

34 Filippakopoulos, P. et al. Selective inhibition of BET bromodomains. Nature 468, 1067–1073 (2010).

35 Ding, J. et al. Effect of miR-34a in regulating steatosis by targeting PPARα expression in nonalcoholic fatty liver disease. Scientific reports 5, 13729 (2015).

36 Herok, M. et al. Chemotherapy of HER2-and MDM2-Enriched Breast Cancer Subtypes Induces Homologous Recombination DNA Repair and Chemoresistance. Cancers 13, 4501 (2021).

37 Chen, Z. S. & Tiwari, A. K. Multidrug resistance proteins (MRPs/ABCCs) in cancer chemotherapy and genetic diseases. The FEBS journal 278, 3226–3245 (2011).

38 Chen, H. H. & Kuo, M. T. Role of glutathione in the regulation of Cisplatin resistance in cancer chemotherapy. Metal-based drugs 2010 (2010).

39 Lehmann, W. et al. ZEB1 turns into a transcriptional activator by interacting with YAP1 in aggressive cancer types. Nature communications 7, 10498 (2016).

40 Boutros, R., Lobjois, V. & Ducommun, B. CDC25 phosphatases in cancer cells: key players? Good targets? Nature Reviews Cancer 7, 495–507 (2007).

41 Hickson, I. et al. Identification and characterization of a novel and specific inhibitor of the ataxia-telangiectasia mutated kinase ATM. Cancer research 64, 9152–9159 (2004).

42 Duan, W. et al. Fanconi anemia repair pathway dysfunction, a potential therapeutic target in lung cancer. Frontiers in oncology 4, 368 (2014).

43 Whittaker, S. R., Mallinger, A., Workman, P. & Clarke, P. A. Inhibitors of cyclin-dependent kinases as cancer therapeutics. Pharmacology & therapeutics 173, 83–105 (2017).

44 Chen, H. H. W. & Macus Tien, K. Role of Glutathione in the Regulation of Cisplatin Resistance in Cancer Chemotherapy. Metal-Based Drugs (2010).

45 Westhoff, G. L., Chen, Y. & Teng, N. N. H. Targeting FOXM1 Improves Cytotoxicity of Paclitaxel and Cisplatinum in Platinum-Resistant Ovarian Cancer. International Journal of Gynecologic Cancer 27, 1602–1609, doi:10.1097/igc.0000000000001063 (2017).

46 Zheng, K. et al. Inhibition of autophagosome-lysosome fusion by ginsenoside Ro via the ESR2-NCF1-ROS pathway sensitizes esophageal cancer cells to 5-fluorouracil-induced cell death via the CHEK1-mediated DNA damage checkpoint. Autophagy 12, 1593–1613, doi:10.1080/15548627.2016.1192751 (2016).

47 Kosinsky, R. L. et al. USP22-dependent HSP90AB1 expression promotes resistance to HSP90 inhibition in mammary and colorectal cancer. Cell Death & Disease 10, 911, doi:10.1038/s41419-019-2141-9 (2019).

48 Yu, J. et al. DNA methyltransferase expression in triple-negative breast cancer predicts sensitivity to decitabine. The Journal of Clinical Investigation 128, 2376–2388, doi:10.1172/JCI97924 (2018).

49 Lang, J.-Y. et al. BikDD Eliminates Breast Cancer Initiating Cells and Synergizes with Lapatinib for Breast Cancer Treatment. Cancer Cell 20, 341–356, 10.1016/j.ccr.2011.07.017 (2011).

50 Serrano-Oviedo, L. et al. Identification of a stemness-related gene panel associated with BET inhibition in triple negative breast cancer. Cellular Oncology 43, 431–444, doi:10.1007/s13402-020-00497-6 (2020).

51 Cheng, Q. et al. Amplification and high-level expression of heat shock protein 90 marks aggressive phenotypes of human epidermal growth factor receptor 2 negative breast cancer. Breast Cancer Research 14, R62, doi:10.1186/bcr3168 (2012).

52 Zhang, Y. et al. miRNA-192-5p impacts the sensitivity of breast cancer cells to doxorubicin via targeting peptidylprolyl isomerase A. The Kaohsiung Journal of Medical Sciences 35, 17–23, 10.1002/kjm2.12004 (2019).

53 Zhang, Z. et al. SRGN crosstalks with YAP to maintain chemoresistance and stemness in breast cancer cells by modulating HDAC2 expression. Theranostics 10, 4290–4307, doi:10.7150/thno.41008 (2020).

54 Herok, M. et al. Chemotherapy of HER2- and MDM2-Enriched Breast Cancer Subtypes Induces Homologous Recombination DNA Repair and Chemoresistance. Cancers 13, 4501 (2021).

55 Gayarre, J. et al. The NER-related gene GTF2H5 predicts survival in high-grade serous ovarian cancer patients. J Gynecol Oncol 27 (2016).

56 Dai, C.-H. et al. RNA interferences targeting the Fanconi anemia/BRCA pathway upstream genes reverse cisplatin resistance in drug-resistant lung cancer cells. Journal of Biomedical Science 22, 77, doi:10.1186/s12929-015-0185-4 (2015).

57 Arora, S., Kothandapani, A., Tillison, K., Kalman-Maltese, V. & Patrick, S. M. Downregulation of XPF–ERCC1 enhances cisplatin efficacy in cancer cells. DNA Repair 9, 745–753, 10.1016/j.dnarep.2010.03.010 (2010).

58 Wang, B. et al. Integrative analysis of pooled CRISPR genetic screens using MAGeCKFlute. 14, 756 (2019).

59 Gao, X. et al. Berberine attenuates XRCC1-mediated base excision repair and sensitizes breast cancer cells to the chemotherapeutic drugs. Journal of Cellular and Molecular Medicine 23, 6797–6804, 10.1111/jcmm.14560 (2019).

60 Liu, M.-m., et al. MiR-30e inhibits tumor growth and chemoresistance via targeting IRS1 in Breast Cancer. Scientific Reports 7, 15929, doi:10.1038/s41598-017-16175-x (2017).

61 Upadhyay, A. K., Ajay, A. K., Singh, S. & Bhat, M. K. Cell Cycle Regulatory Protein 5 (Cdk5) is a Novel Downstream Target of ERK in Carboplatin Induced Death of Breast Cancer Cells (Supplementary Data). Current Cancer Drug Targets 8, 741–752, doi:10.2174/156800908786733405 (2008).

62 Syu, J.-P., Chi, J.-T. & Kung, H.-N. Nrf2 is the key to chemotherapy resistance in MCF7 breast cancer cells under hypoxia. Oncotarget 7, 14659–14672, doi:10.18632/oncotarget.7406 (2016).

63 Chen, Z., Huang, J., Feng, Y., Li, Z. & Jiang, Y. Profiling of specific long non-coding RNA signatures identifies ST8SIA6-AS1 AS a novel target for breast cancer. The Journal of Gene Medicine 23, e3286, 10.1002/jgm.3286 (2021).

64 Lu, H. et al. Chemotherapy triggers HIF-1–dependent glutathione synthesis and copper chelation that induces the breast cancer stem cell phenotype. Proceedings of the National Academy of Sciences 112, E4600–E4609, doi:doi:10.1073/pnas.1513433112 (2015).

65 Nedeljković, M., Tanić, N., Prvanović, M., Milovanović, Z. & Tanić, N. Friend or foe: ABCG2, ABCC1 and ABCB1 expression in triple-negative breast cancer. Breast Cancer 28, 727–736, doi:10.1007/s12282-020-01210-z (2021).

66 Mohammadi, A. et al. The Urtica dioica extract enhances sensitivity of paclitaxel drug to MDA-MB-468 breast cancer cells. Biomedicine & Pharmacotherapy 83, 835–842, 10.1016/j.biopha.2016.07.056 (2016).

67 Deng, X. et al. Combined Phosphoproteomics and Bioinformatics Strategy in Deciphering Drug Resistant Related Pathways in Triple Negative Breast Cancer. International Journal of Proteomics 2014, 390781, doi:10.1155/2014/390781 (2014).

68 Dou, D. et al. CircUBE2D2 (hsa_circ_0005728) promotes cell proliferation, metastasis and chemoresistance in triple-negative breast cancer by regulating miR-512-3p/CDCA3 axis. Cancer Cell International 20, 454, doi:10.1186/s12935-020-01547-7 (2020).

69 Westhoff, G. L., Chen, Y. & Teng, N. N. H. Targeting Foxm1 Improves Cytotoxicity of Paclitaxel and Cisplatinum in Platinum-Resistant Ovarian Cancer. International Journal of Gynecologic Cancer 27, 887–894, doi:10.1097/igc.0000000000000969 (2017).

70 Zhang, Y. et al. Elevated Aurora B expression contributes to chemoresistance and poor prognosis in breast cancer. Int J Clin Exp Pathol 8, 751–757 (2015).

